# PVN microglia via P2Y_12_ transmit hemodynamic signal to promote sympathetic excitation in hypertension

**DOI:** 10.1101/2023.08.29.555298

**Authors:** Bo Wei, Guo Cheng, Li Li, Qihang Sun, Qianqian Bi, Cheng Lu, Chunyou Yin, Ningting Chen, Miner Hu, Haoran Lu, Zhechun Hu, Genxiang Mao, Yan Gu, Shu Wan, Xiaoli Liu, Xiao Z. Shen, Peng Shi

## Abstract

Hypertension is usually accompanied with an elevated sympathetic tonicity, but how sympathetic hyperactivity is triggered is not fully understood. Recent advances reveal that microglia-centered neuroinflammation contributes to sympathetic excitation in hypertension. In this study, we performed a temporospatial analysis of microglia at both morphological and transcriptomic levels, and found that microglia in the hypothalamic paraventricular nucleus (PVN) were early responders to hypertensive challenges. PVN is the central hub for maintaining cardiovascular function via regulation of fluid balance and sympathetic outflow. Comprehensive vasculature analyses unveiled that PVN was characterized by high capillary density, thin vessel diameter, and complex vascular topology among brain regions. As such, PVN is susceptible to the penetration of ATP released from the vasculature in response to hemodynamic disturbance after blood pressure increase. ATP ligation to microglial P2Y_12_ receptor is responsible for the microglial accumulation and activation in the PVN. Furthermore, either pharmacological blockade or genetic ablation of microglial P2Y_12_ could substantially restrain blood pressure increase under hypertensive challenge. Together, these findings disclose that a unique vasculature pattern results in the vulnerability of PVN pre-sympathetic neurons to hypertension-associated insults, which is mediated by microglia.

## INTRODUCTION

Essential hypertension is defined by a rise of blood pressure (BP) with unknown etiology, which is a significant portion of hypertension, affecting ∼90% hypertensive patients^1^. Although its etiology is multifactorial, a significant portion of essential hypertension have two common features: elevated sympathetic nerve activity (sympathoexcitation) and persistent systemic inflammation^2–8^. Elevated sympathetic nerve activity has been reported in ∼50% of untreated essential hypertensive patients^9–11^. It has been demonstrated that patients with borderline hypertension who later proceeded to hypertension was due to a prolonged sympathoexcitation^12–14^. Moreover, people with cardiovascular hyperreactivity to mental stress was predicted to develop hypertension with elevated sympathetic activity^15, 16^, further highlighting the association between essential hypertension and sympathetic over-activation. On the other hand, more and more evidence showing development of inflammation secondary to hypertension emerges in humans^17–20^. These two factors are actually interwoven in the central nervous system (CNS), as neuroinflammation, especially that occurring in the hypothalamic paraventricular nucleus (PVN), flares a sustained sympathoexcitation, thus further contributing to BP increase^21–23^. Microglia are the dominant immune cells in the parenchyma of the CNS. Many studies including ours unravel that microglia could directly modulate neuronal activities via various mechanisms in different brain regions^24–29^. Indeed, microglia activation constitutes a pivotal component of the neuroinflammation associated with hypertension^4, 6^. Consistently, either central anti-inflammation treatments or microglia depletion could remarkably attenuate hypertension progression^23, 30^. However, the fundamental mechanism underlying microglial activation in the PVN during hypertension development still remains unidentified.

PVN is located in the hypothalamus bilaterally adjacent to the third ventricle^31–33^. Despite its small size, PVN is specifically devoted to maintaining autonomic and neuroendocrine homeostasis, which is highly conserved from zebrafish to humans^34–37^. PVN consists of magnocellular and parvocellular neurons. The magnocellular neurons project mainly to the posterior pituitary gland to secrete hormones including arginine vasopressin (AVP) which regulates body fluid balance^38^; the parvocellular neurons are primarily composed of two types of cells which project to the sympathetic vasomotor neurons in the brainstem rostroventrolateral medulla (RVLM)^5, 39^, and to the anterior pituitary as the starting point of the hypothalamic-pituitary-adrenal axis^40^, respectively. Recent studies indicate that PVN neurons, in particular those involved in AVP release and sympathetic activity, would be activated in the context of local hypoxia^41, 42^. As such, they could respond in a timely manner to insufficient blood perfusion in occasions like hypovolemic shock, leading to a collective effect of compensatory systemic BP increase which is critical for survival. However, little is known about whether there are anatomical specificities in the PVN attributable to a quick response by PVN to hemodynamic disturbance.

In the present study, we show that the PVN is characterized by high capillary density and complex vascular topology relative to most other brain regions by high resolution microscopy. This topological feature of the vasculature predisposed the PVN to the vasculature-derived ATP owing to the BP increase. Here, the elevated level of environmental ATP stimulated microglia via purinergic P2Y_12_ receptor, leading to microglial accumulation in the PVN and activation. Deprivation of microglial P2Y_12_ signaling by pharmacological or genetic approach could restrain sympathetic overflow and BP increase during hypertensive challenge. Thus, the PVN vasculature exemplifies an anatomical structure which is advantageous for survival in wild environment turning disadvantageous in a prevalent disease of modern lifestyle.

## RESULTS

### PVN microglia are early responders to BP increase

Previous studies demonstrate that in a variety of murine hypertension models, there is reactive microgliosis^43^ characterized by increased microglial density and microglial activation, particularly in the PVN^23, 44, 45^. However, this reaction of microglia has only been reported in the chronic established stage of hypertension, i.e., 14-28 days post hypertensive induction. To investigate the dynamics and regional features of microgliosis in response to hypertension, we examined microglia in the hypothalamic PVN and brainstem RVLM, two sympathetic-regulatory nuclei, and the somatosensory barrel cortex (S1BF) which served as an autonomic function-unrelated control region in the acute BP climbing stage of hypertension, i.e., 1-3 days post hypertensive induction (Extended Data Figure 1a)^30^. In resting state, the densities of microglia were comparable between these brain regions (Figure 1a). However, PVN displayed a significant increase of microglial density as early as 3 days after mice were treated with N (ω)-nitro-L-arginine methyl ester (L-NAME), a vasodilation inhibitor widely used to induce hypertension in animals (Figure 1a). In contrast, both RVLM and SIBF displayed normal microglia distribution at this time point. Angiotensin II (Ang II) has a distinctive mechanism from L-NAME in inducing BP increase. When we treated mice with Ang II, we also observed an increase of microglia number exclusively in the PVN on day 3 post induction (Figure 1a), supporting the idea that an acute microglial accumulation in the PVN is a hypertension-elicited response. An increased microglia density was still evident on day 14 post hypertension induction, consistent with previous reports (Extended Data Figure 1b)^30, 46–48^. In addition to increased density, microglia in the PVN displayed greater morphological changes relative to their counterparts in the other regions on day 3 post hypertension induction (Figure 1b). PVN microglia exhibited enlarged soma volume and decreased dendrite complexity manifested by Sholl analysis, further confirming a microgliosis phenotype specific to the PVN during the early stage of hypertension.

**Figure 1.**
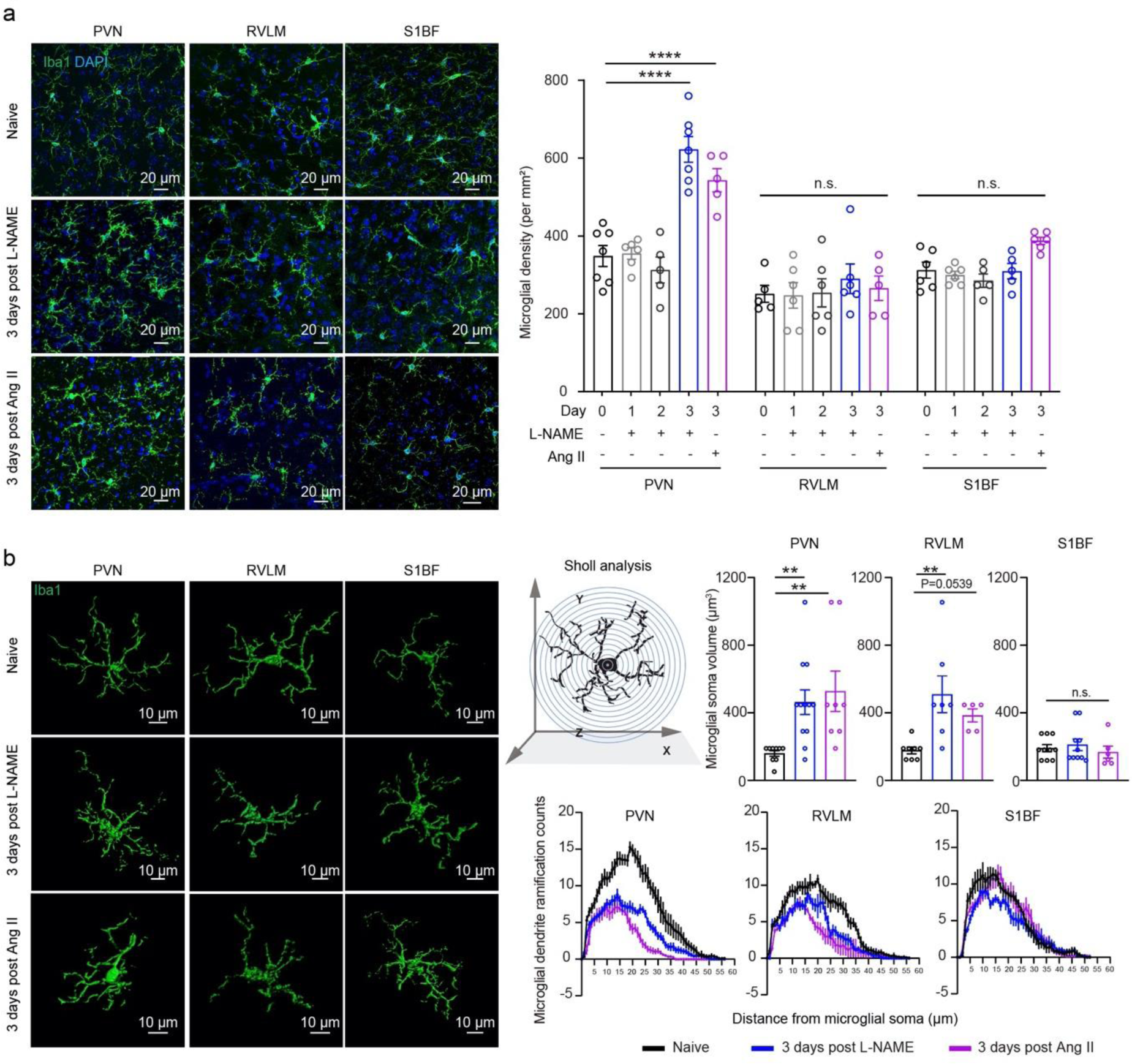
Density and morphological changes of microglia in different nuclei in the early stage of hypertension. **a**, Representative confocal images and density analysis of Iba1^+^ microglia in the PVN, RVLM and somatosensory barrel cortex (S1BF) from normotensive controls (Day 0), and mice after 1 to 3 days of either L-NAME-or Ang II treatment. Each dot indicates the average of four 160 ×160 μm^2^ fields of view (FOVs) in the indicated region of one mouse. **b**, Representative 3D-reconstructed Iba1^+^ microglia from (a). Quantification of microglial soma volume and branch ramification complexity (illustration of Sholl analysis). Each dot indicates the average of 10-15 microglia in the indicated region of one mouse. n.s., not significant. ** P<0.01, **** P<0.0001 by one way ANOVA with Tukey’s post hoc test.

To further investigate hypertension-elicited microglial alteration, we analyzed the transcriptomes of microglia derived from the hypothalamus (containing PVN), brain stem (containing RVLM), and cerebrocortex in resting state and on day 3 post L-NAME or Ang II treatment. Microglia (the CD11b^+^CD45^low^ population) from the abovementioned regions were individually sorted, followed by bulk RNA sequencing (RNA-seq) (Figure 2a). The RNA-seq data indicated that the signature genes relevant to microglia ontology and functions, *e.g.*, *Hexb*, *Csf1r*, *Cx3cr1*, *Selplg*, *P2ry12*, were highly expressed in all the samples examined; in contrast, characteristic genes for neurons, astrocytes, oligodendrocytes, or endothelial cells were barely detected, validating their microglia identity (Extended Data Figure 2a). For bulk RNA-seq, we obtained comparable microglial counts from hypothalamus (2,169±855) and brain stem (2,125±852) yielded from the similar tissue size (2×2×2 mm^3^); and 10,012±12 microglia from the entire cortex of the same batches of animals. Although the cell counts were not exactly the same between the cortex and the other regions, the detected gene numbers were very similar (Extended Data Figure 2b), which was also confirmed by Pearson correlation when comparing the normalized expression of genes (R > 0.98) (Extended Data Figure 2c). Consistent with recent reports, microglia displayed limited regional divergence in resting state^49–51^. Intriguingly, hierarchical clustering analysis revealed that the replicates of PVN tended to cluster together in both hypertension models (Extended Data Figure 2d). When the differentially expressed genes (DEGs) (Log_2_ Fold Change [FC]>1; adjusted P_value_< 0.05) between hypertensive and normotensive samples from the same region were compared, microglia derived from different models displayed differential profiles (Extended Data Figure 2e). We reasoned that the common DEGs altered in both hypertension models unambiguously reflect the pressor effects. Thus, we focused on the common DEGs. This yielded smaller DEG subsets; and hypothalamic microglia outnumbered their counterparts derived from the other two regions in upregulated common DEGs (33 vs. 5 in the brain stem vs. 20 in the cortex) (Figure 2b, these DEGs are listed in Extended Data Figure 2f). Moreover, David Gene ontology (GO) analysis based on the common DEGs unveiled region-specificity of microglia at the early stage of hypertension as the GO terms of hypothalamic microglia displayed smaller P values compared to the ones derived from the other two brain regions (Figure 2c). And as indicated by the GO terms, hypothalamic microglia were particularly strengthened in the function of reactive oxygen species, inflammation and migration after hypertension induction; in contrast, brainstem and cortical microglia were both characterized by altered metabolic and biosynthesis processes, indicating a shift to pro-inflammatory phenotype only occurring in the hypothalamic microglia. Collectively, these data demonstrate a regional specificity of microglial perturbation upon pressor challenge at the early stage of hypertension, as PVN-residing microglia were prominently affected in the brain.

**Figure 2.**
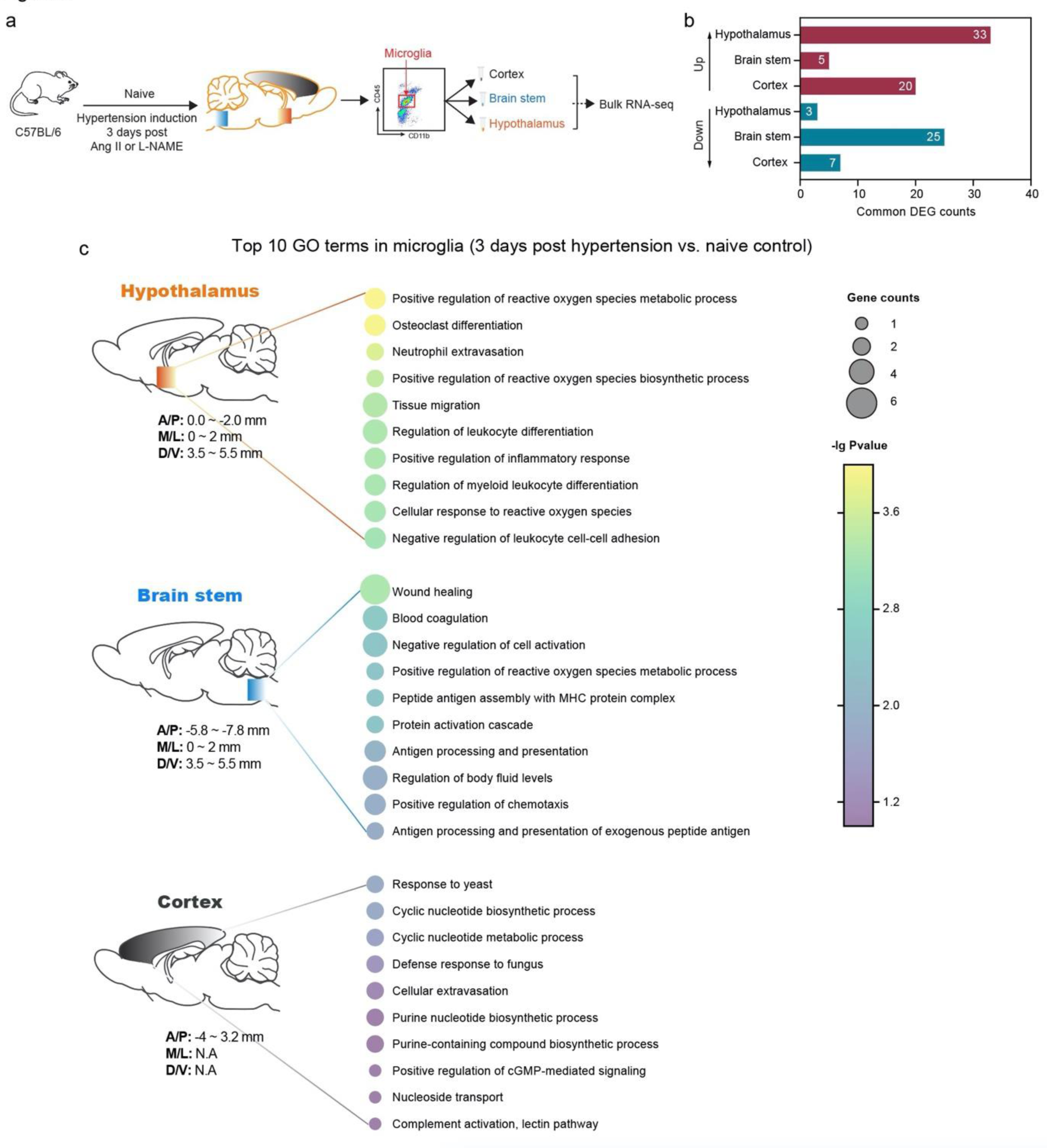
Transcriptional changes of microglia from different nuclei in the early stage of hypertension. **a**, Schematic outline for transcriptome analysis of dissociated microglia from hypothalamus, brain stem and cortex from the same batch of animals. **b**, Numbers of common differentiated expression genes (DEGs) from microglia of the indicated brain regions 3 days post Ang II or L-NAME treatment (in comparison to the microglia dissociated from the same regions of naïve controls). **c**, Top 10 gene ontology (GO) annotations based on P values for the common DEGs of microglia derived from the indicated regions.

It is well established that the animal hypertension models induced by L-NAME or Ang II, as well as many patients with essential hypertension, have a sympathetic component^52, 53^. To understand the temporal relationship between microglial activation and sympathetic excitation along hypertension progress, we analyzed the PVN pre-sympathetic neuronal activity by electrophysiological recording. PVN pre-sympathetic neurons were pre-labeled by using the retrograde fluorescent tracer from the downstream RVLM^24^; two weeks later, mice were treated with L-NAME for hypertension induction.

We found that PVN pre-sympathetic neurons displayed significantly elevated resting membrane potential and evoked action potential as early as 5 days post L-NAME (Extended Data Figures 3a-b). Consistently, a systemic sympathetic excitation, as valued by plasma norepinephrine (NE), was also significantly increased on Day 5 (Extended Data Figure 3c), accompanied by an elevation of NE level in multiple organs (Extended Data Figure 3d). These data indicate that in the initial climbing stage of hypertension, microglial activation precedes a systemic sympathetic activation.

### Increased extracellular ATP is responsible for microglial accumulation in the PVN in the early stage of hypertension

Both Ki67 staining and BrdU incorporation assay showed negligible proliferating microglia in the PVN on day 3 post L-NAME treatment (Extended Data Figure 4a-b), suggesting that migration rather than proliferation was responsible for microglial accumulation in the PVN in the early stage of hypertension, which is consistent with the bulk RNA sequencing result (Figure 2c). To understand the mechanism underlying microglial migration to the PVN, we first investigated chemokine receptor CX3CR1 which is highly expressed in microglia and is important for the proper migration of microglia to sites of injury^54, 55^. CX3CR1-deficient *Cx3cr1*^GFP/GFP^ mice, which have a GFP sequence replacing the coding exon of the *Cx3cr1* gene, was used to investigate a role of CX3CR1. *Cx3cr1*^GFP/GFP^ mice developed a normal pressor response to L-NAME (Extended Data Figure 5a) and displayed a comparable increase of PVN microglia (Extended Data Figure 5b) to their wild-type littermates 3 days post hypertension induction, arguing against a role of CX3CR1.

Considering that hemodynamic disturbance is the most prominent alteration associated with BP increase at the early stage of hypertension, we hypothesized that a vasculature-derived molecule(s) is responsible for microglia accumulation in the PVN. We recently demonstrated that ATP release from red blood cells and blood vessel is one of early events following BP increase^56^, possibly due to an increased shear stress in the blood flow^57^. Moreover, the elevated extracellular ATP instigates a systemic inflammation associated with hypertension^56^. Indeed, ATP is a potent attractant for microglia migration in the CNS via purinergic receptors^58–60^. Therefore, we next investigated environmental ATP in the periphery and in the brain parenchyma after hypertension. A significant rise of plasma ATP was observed starting on day 1∼2 post hypertension induction (Figures 3a-b), predating microglial accumulation in the PVN. Co-treatment of hydralazine, a vasodilator, could restore both BP and plasma ATP to normal levels (Figures 3a-b), confirming an effect of BP increase on the elevation of plasma ATP. To investigate whether an increase of microenvironmental ATP is the cause of microglia accumulation in the PVN, we intracerebroventricularly (ICV) infused the mice with apyrase, an ATP hydrolase, to promote ATP degradation in the brain interstitium (Figure 3c). Supporting our hypothesis, removal of extracellular ATP could almost completely abrogate the increase of PVN microglia in the early stage of hypertension (Figure 3d). Moreover, mice with apyrase infusion displayed a restrained increment of BP in comparison with mice ICV treated with vehicle in response to either L-NAME or Ang II (Figure 3e). These data indicate that an elevation of environmental ATP is a contributing factor of microglia accumulation in the PVN, which further contributes to BP increase.

**Figure 3.**
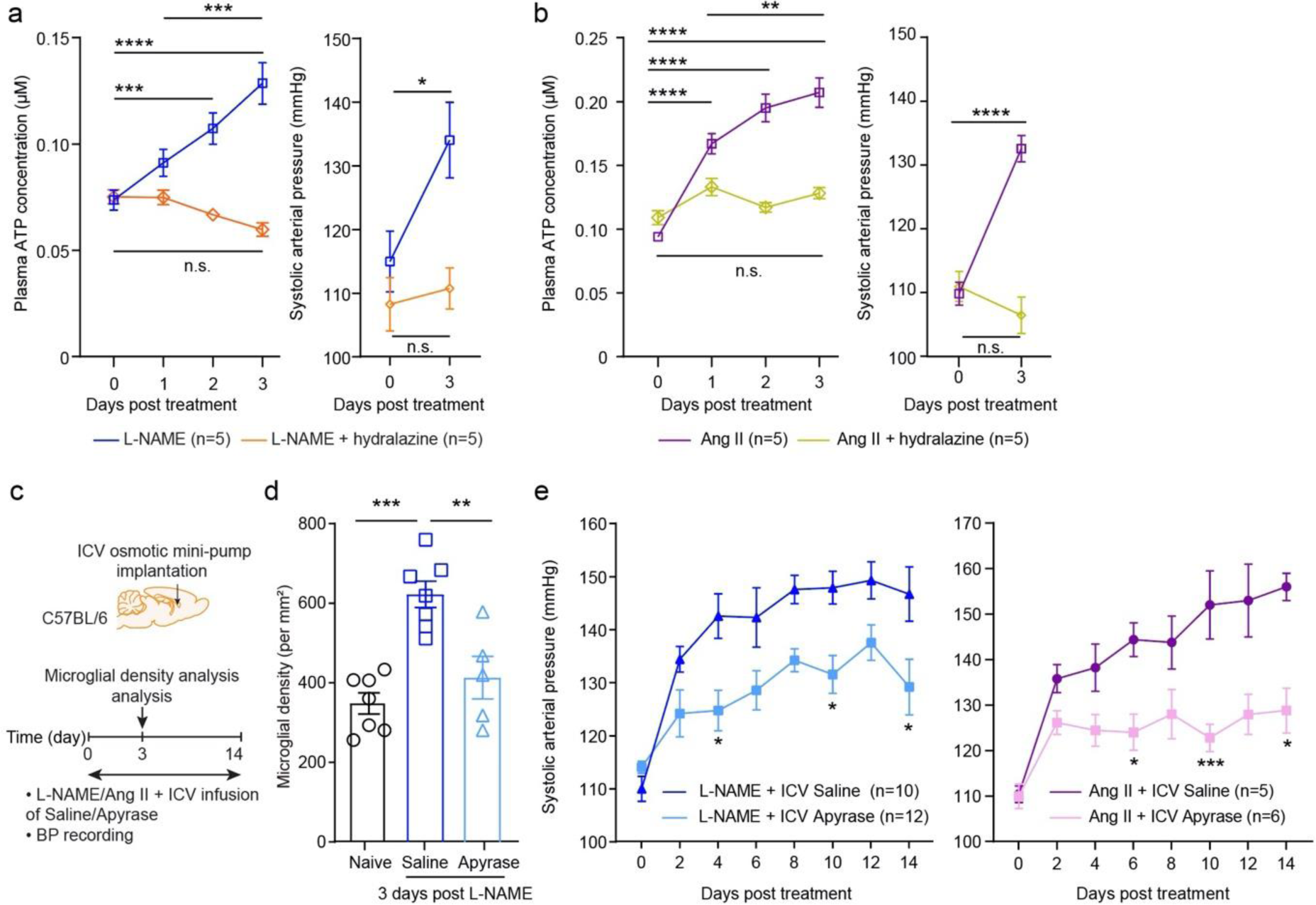
Degrading environmental ATP in the brain suppresses microglial recruitment in the PVN and BP increase in hypertension. **a** and **b**, C57BL/6 mice were treated with L-NAME (a) or Ang II (b), and some of them had a co-treatment of hydralazine. Blood plasma ATP was quantified (left panel) and BP was recorded (right panel) as well in the indicated time points. **c**, Schematic outline of experimental protocol with apyrase ICV infusion for experiments in (d) and (e). **d**, Quantification of microglial density in the PVN 3 days post L-NAME induction. **e**, Systolic arterial pressure was measured during the hypertensive challenge. * P<0.05, ** P<0.01, *** P<0.001 and **** P<0.0001 by one-way ANOVA with Tukey’s post hoc test (d); and two-way ANOVA with Tukey’s post hoc test (a, b, e).

### Vascular architecture makes PVN vulnerable to blood-derived ATP

To investigate whether the disparity in microglial accumulation between brain regions during the early stage of hypertension is due to a difference in local ATP levels, we next adopted a recently developed ATP sensor designed for detecting extracellular ATP^61, 62^. To this end, we simultaneously transfected the PVN and the medial prefrontal cortex (mPFC) which served as a deep control region of the same mice with an adenovirus vector. This vector expresses the probe GRAB_ATP1.0_ which contains an extracellular motif of human P2Y_1_ receptor as the extracellular ATP-binding scaffold linked at its N and C termini with circularly permuted enhanced GFP moieties. ATP binding induce a conformational change of the probe, resulting in proximity of the GFP moieties which gives off fluorescence under an excitation wavelength ∼500 nm. Two optical fibers were positioned in the transfected regions on the same day of viral injection, followed by a 2-week recovery. On the recording day, we mounted the detection system onto the optical fibers, and recorded real-time ATP signal in the PVN and cortex of the same mouse simultaneously (Figure 4a). Since the procedure of mounting the detection system would unavoidably bring in varying background noise, a comparison between the values of ATP signals acquired from the same region of the same mouse in different days or acquired from different fibers (brain regions) was impractical. Therefore, we adopted an alternative strategy by comparing the changes of the ATP signals collected from different regions of the same mouse before and after a bolus injection of Ang II (Figure 4a). In a preliminary experiment, we observed that an injection of Ang II (7.2 mg/kg body weight) could acutely raise BP by ∼40 mmHg in ∼2 min, and this pressor effect could last for over 20 min (Extended Data Figure 6a). Concomitantly, plasma ATP level would significantly rise post Ang II treatment (Extended Data Figure 6b left panel). However, same volume of saline injection did not alter BP or plasma ATP levels (Extended Data Figure 6b right panel). We observed that the pressor reaction induced by Ang II generated an increment of ATP signal in both the PVN and cortex at 2 min post Ang II administration (Figure 4b). However, there was a greater rise of ATP signals in the PVN than the cortex. ATP has a molecular weight of 507 Da, right within the cut-off molecular size for BBB penetration (400∼600 Da)^63^. To interrogate whether ATP is indeed prone to accumulate in the PVN over other brain regions, we intravenously infused rhodamine B, an amphiphilic fluorescent probe with molecular weight (479 Da) close to ATP, and measured its penetration into the parenchyma of 3 disparate brain regions including hypothalamus where PVN locates. It showed that while rhodamine B could be detected in all the tested regions, hypothalamus had the highest permeability, implying a greater dissemination of blood vessel-borne small molecules in the local (Figure 4c).

**Figure 4.**
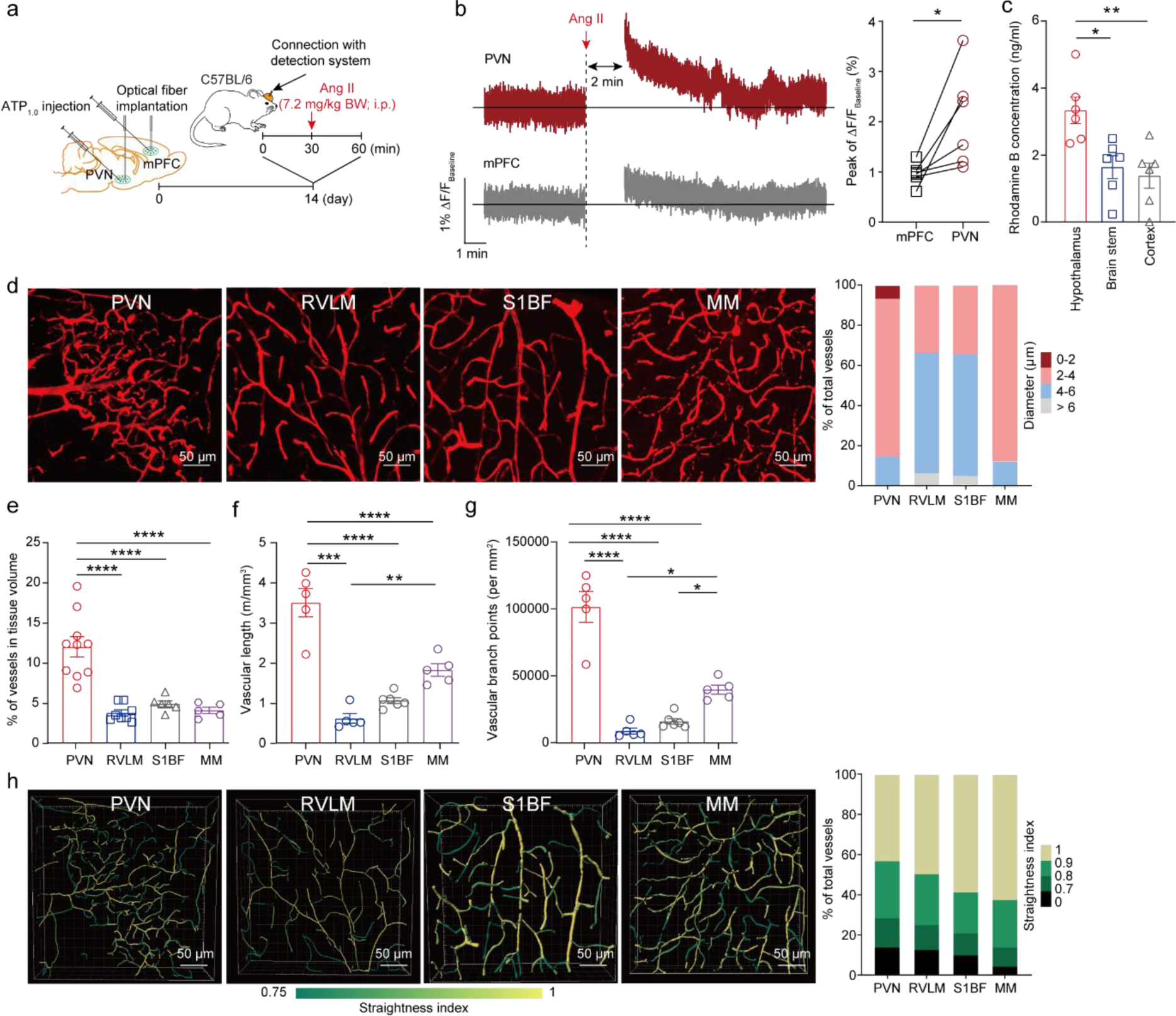
*In vivo* quantification of environmental ATP in the brain parenchyma and the profiles of cerebral vasculatures in different nuclei. **a**, Schematic outline of transfecting an AAV-mediated ATP sensor (ATP1.0) in the cortex (mPFC) and PVN, followed by optical fiber implantation for *in vivo* imaging of ATP signals before and after a bolus i.p. injection of Ang II. **b**, Representative fluorescence intensity changes of ATP1.0 and the statistical comparison of peak ΔF/F_Baseline_ measured in the PVN and cortex before and after i.p. injection of Ang II. **c**, Fifteen minutes after i.v. infusion of rhodamine B, mice were transcardiacally perfused with PBS. Fluorescence intensities of rhodamine B in the parenchyma of the indicated regions were quantitated. **d**, Representative images of cerebral vasculature illustrated by fluorophore-tagged dextran (150 kDa) in the PVN, RVLM, S1BF and MM, and the quantitation of vessels with different diameters is shown. The percentages are the averages from 6-8 mice. **e**-**g**, Quantitative profiling of vascular volume (e), length (f) and degree of ramification (g) in the indicated nuclei. Each dot indicates the average of four 319 ×319 μm^2^ FOVs in the indicated region of one mouse. **h**, Representative annotated images and quantification for vascular straightness in the indicated nuclei. The percentages are the averages from 6-8 mice. * P<0.05, ** P<0.01, *** P<0.001, **** P<0.0001 by paired t-test in (b); and by one-way ANOVA with Tukey’s post hoc tests in (c, e-g).

Identifying ATP as the culprit causing microglia recruitment in the PVN could not resolve the puzzle of a spatial specificity in ATP diffusion favoring PVN in the brain. Especially, there was no detectable BBB damage in the PVN 3 days post hypertension induction (Extended Data Figure 6c). We hence speculated that the PVN possesses a unique vascular arrangement predisposing this nucleus to the fallouts of hemodynamic disturbance, such as ATP release. To test this hypothesis, we performed quantitative analyses of the vascular network across brain regions. Intriguingly, it showed that the majority (85.3 ± 5.6%) of the blood vessels in the PVN were ultra-small (< 4 μm in diameter), while the vessels in this extremely small size only constituted 33.9 ± 35.2% and 36.1 ± 20.7% of the blood vessels in the RVLM and S1BF, respectively (Figures 4d). These ultra-small vessels were scarce in the expression of α-smooth muscle actin (α-SMA) but enriched in CD13 expression, confirming their capillary identity^64^ (Extended Data Figure 7). Moreover, analysis of coronary sections with a 70 μm Z-stack unveiled that the PVN possessed the largest vascular volume and highest vascular length in comparison to the RVLM and S1BF (Figure 4e-f). These observations are consistent with a recent study of large-scale cerebral vascular reconstructions which also revealed that PVN was one of the nuclei possessing the highest degree of vessel density^65^. Different from our intravascular labeling strategy, vasculature was outlined by immunolabeling of vessel wall in that study. That study also showed that median mammillary nucleus (MM) which also locates in the hypothalamus had the highest vessel length in the brain. We thus also incorporated MM for a parallel comparison in our study. Surprisingly, we found that PVN was even greater in both vascular volume and length than MM (Figure 4e-f). To unveil the characteristics of PVN vasculature in more details, we further investigated the regional variations in the topological organization of brain vasculature. To this end, we first identified all branch points at which vessels split or join. It showed that the PVN had the most branch points compared to the other tested regions (Figure 4g). Moreover, three-dimension (3D) reconstruction and filament module analysis revealed that the vasculature in the PVN possessed the lowest degree of straightness among the brain regions examined (Figure 4h). Altogether, the PVN has a greater capillary density with a smaller vascular diameter, higher ramification and less straightness than those most of other brain regions. As smaller diameter and higher ramification of capillary, together with higher capillary density, are associated with slower flow velocity and higher exchange rate^66–68^, these features warrant a more sufficient dispersion of ATP from the blood into the parenchyma in the PVN relative to other brain regions during hypertension.

### Disruption of ATP-P2Y_12_ signaling abolishes microglial accumulation and activation in the PVN, and alleviates pressor response

Purinergic P2Y_12_ receptor is exclusively expressed in the microglia among all CNS cell populations; also, it is a signature marker distinguishing microglia from peripheral myeloid cells^69^. Importantly, it has been shown that P2Y_12_ mediates microglia migration to focal BBB lesion^58^. We thus hypothesized that the ATP-P2Y_12_ engagement mediated the microgliosis in the PVN under hypertension. To explore this, we employed both pharmacological and genetic approaches. ICV infusion of PSB-0739 (Figure 5a), a selective antagonist of P2Y_12_ receptor^28^, could not only significantly reduce microglial accumulation in the PVN (Figure 5b), but also markedly alleviated BP increase after hypertension induction (Figure 5c). To further investigate the role of P2Y_12_, we crossed *Cx3cr1*^CreERT2/+^ mice with *P2ry12*^fl/fl^ mice; the offspring (hereafter termed as *P2ry12*^MG-KO^) could be ablated with microglial P2Y_12_ expression upon tamoxifen treatment (Figure 5d). Indeed, there was 68% reduction of *P2ry12* transcripts in the whole brain of *P2ry12*^MG-KO^ mice 3 weeks after tamoxifen treatment in reference to the *P2ry12*^fl/fl^ mice (Extended Data Figure 8a). Moreover, we analyzed the efficiency of P2Y_12_ abrogation by flow cytometry analysis, which revealed a marked reduction of the ratio of P2Y_12_ ^+^ microglia (from 81.6±6.0% in control brain to 17.9±7.1% after tamoxifen treatment), as well as a remarkable decline in microglial P2Y_12_ mean fluorescence intensity (MFI) (7,798±3,978 a.u. vs. 49,836±14,058 a.u.) (Extended Data Figure 8b), verifying the faithfulness of this genetic approach. Consistent with a previous report^70^, P2Y_12_ deficiency did not alter microglial distribution across brain in resting state (Extended Data Figure 8c). However, abrogation of microglial P2Y_12_ expression abolished microglia number increase in the early stage of hypertension (Figure 5e). More importantly, deprivation of microglial P2Y_12_ expression significantly suppressed BP increase along either L-NAME or Ang II treatment (Figure 5f).

**Figure 5.**
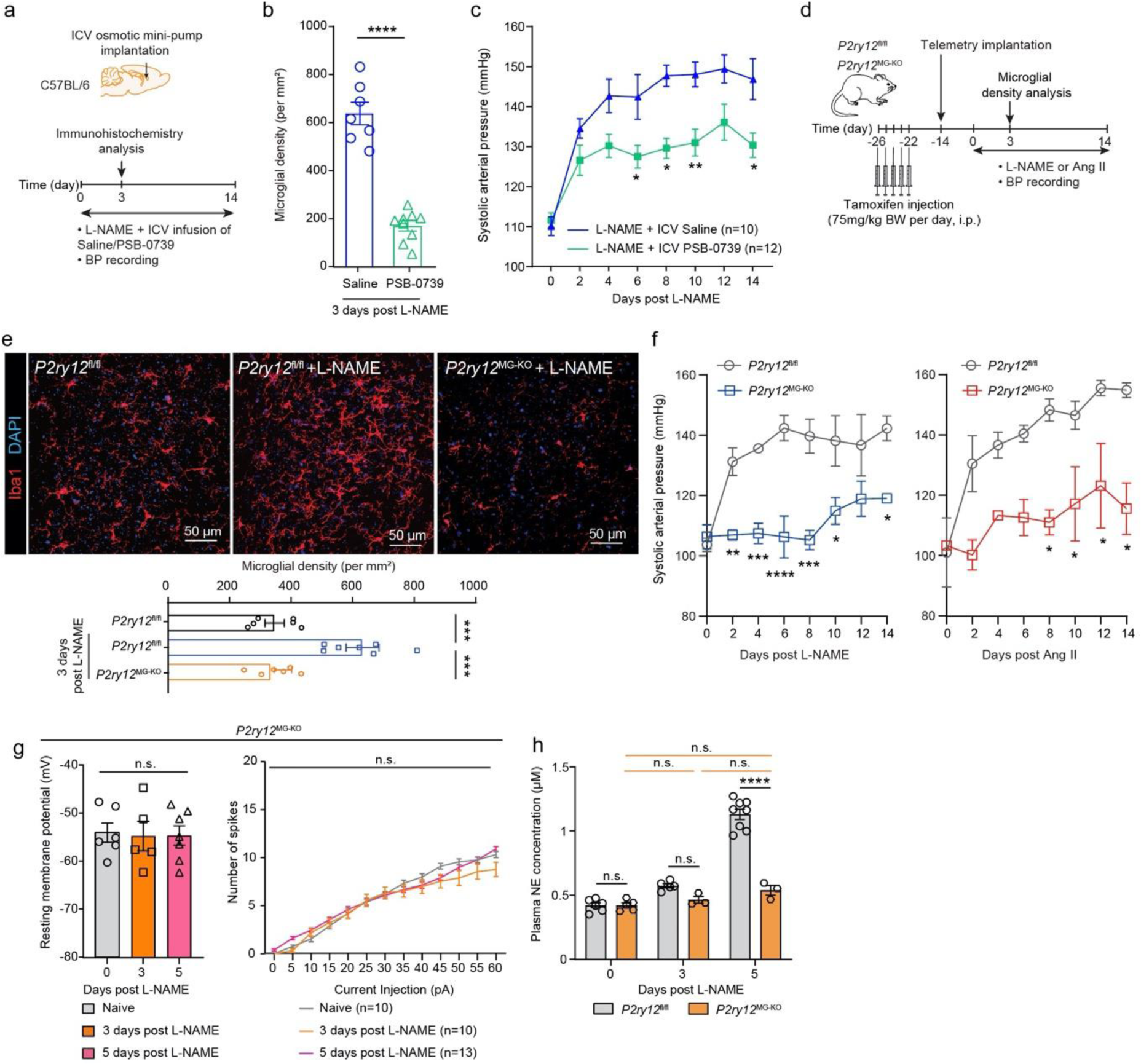
Ablation of P2Y_12_ signaling attenuates L-NAME-elicited pressor response and sympathetic tonicity. **a**, Schematic outline for the Experiments in (b) and (c). ICV infusion of saline or P2Y_12_ antagonist PSB-0739 by minipumps to C57BL/6 mice during L-NAME hypertensive challenge. **b**, Density of Iba1^+^ microglia in the PVN with the indicated ICV treatments. **c**, Systolic arterial pressure during the indicated ICV treatments. **d**, Schematic outline for the experiments shown in (e-f) using the *P2ry12*^fl/fl^ and *P2ry12*^MG-KO^ mice. **e**, Representative images and quantitation of Iba1^+^ microglia in the PVN of the indicated mice with or without 3-day L-NAME treatment. **f**, Systolic arterial pressure responses to either L-NAME or Ang II in *P2ry12*^fl/fl^ and *P2ry12*^MG-KO^ mice. n = 4. **g**, Resting membrane potential, membrane resistance and evoked action potential of PVN pre-sympathetic neurons in naïve *P2ry12*^MG-KO^ mice or the mice after 3-day or 5-day L-NAME treatment. **h**, Plasma norepinephrine (NE) levels in the indicated time points during L-NAME treatment. In (b) and (e), each dot indicates the average of four 160 ×160 μm^2^ FOVs in the PVN of one mouse. n.s., not significant, * P<0.05, ** P<0.01, *** P<0.001, and **** P<0.0001 by unpaired t-test in (b), one-way ANOVA with Tukey’s post hoc analysis (e) and (g left panel), two-way ANOVA with Tukey’s post hoc analysis in (c), (f), (g right panel) and (h).

To further substantiate that the abolished pressor response is microglial P2Y_12_-depleted mice was due to a suppressed activity of PVN pre-sympathetic neurons, we analyzed the electrophysiology of PVN-RVLM neurons. Indeed, *P2ry12*^MG-KO^ mice showed no statistical difference in resting membrane potential and evoked action potential in the PVN-RVLM neurons from control mice after hypertension induction (Figure 5g). Consistently, *P2ry12*^MG-KO^ mice exhibited no increase of plasma NE during the multiple-time observation (Figure 5h).

### C/EBPβ mediates ATP-P2Y_12_ signaling in microglial activation in hypertension

In addition to migration, P2Y_12_ signaling has been implicated in regulating microglial activation^59, 71^. To interrogate whether P2Y_12_ was involved in the activation of microglia in the PVN after hypertension, we resorted to the RNA-seq data. Among the 33 common upregulated early DEG in the PVN microglia under both L-NAME and Ang II treatment was *Cebpb* (Extended Data Figure 2f and Figure 6a), the gene encoding C/EBPβ which is a master transcription factor orchestrating microglial activation in both brain injury and neurodegenerative diseases^72–74^. Importantly, it was reported that suppressing C/EBPβ expression in microglia could dampen neuroinflammation^75^. Indeed, Kyoto Encyclopedia of Genes and Genomes (KEGG) pathway analysis identified 6 genes of the common upregulated DEGs which were positively regulated by P2Y_12_ signaling, and *Cebpb* was the only direct downstream gene of P2Y_12_ among them (Figure 6b). To be noted, *Cebpb* was a component of most of the top GO terms (Figure 6c), especially those associated with leukocyte migration, differentiation and inflammatory responses.

**Figure 6.**
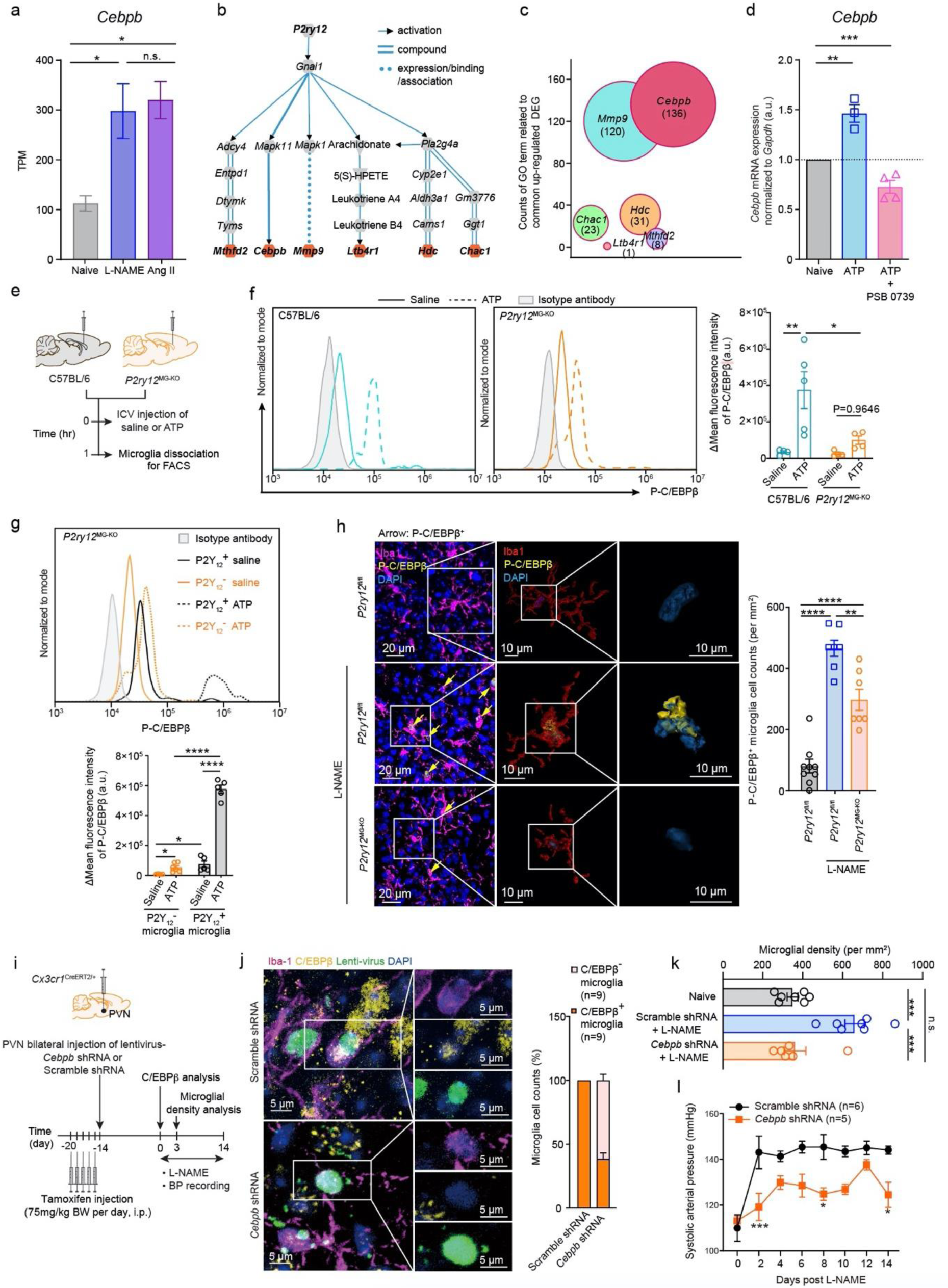
P2Y_12_ activates C/EBPβ in the PVN microglia during hypertension. **a**, Bulk RNA-seq data showed that *Cebpb* transcripts in the microglia dissociated from PVN in naïve C57BL/6 mice or their littermates subjected to either L-NAME or Ang II treatment for 3 days. **b**, KEGG pathway analysis showed the projected links between P2Y_12_ signaling and 6 genes (highlighted in orange) among the up-regulated common DEGs in hypertensive vs. normotensive mice. **c**, Numbers of GO terms comprising the above 6 downstream DEGs of P2Y_12_ signaling. **d**, *Cebpb* mRNA expression in microglia BV2 cells with or without 1-hr ATP treatment. One group of cells had co-treatment of ATP and P2Y_12_ antagonist PSB-0739. **e**, Schematic outline of experiment for (f): C57BL/6 and *P2ry12*^MG-KO^ mice received ICV injection of either saline or ATP treatment, and one hour later, brains were dissociated from above fours cohorts for flow cytometry analysis. **f**, Representative histograms and statistical analyses of phosphorylated C/EBPβ (P-C/EBPβ) in CD11b^+^CD45^low^ microglia at 1 hr post the indicated treatments in above two cohorts of mice. Data was presented as the difference in mean fluorescent intensity (Δ) between P-C/EBPβ staining and isotype antibody staining. **g**, Representative histogram and statistical analysis of P-C/EBPβ in P2Y_12_^+^ and P2Y_12_^−^microglia at 1 hr post the indicated treatments in the tamoxifen-treated *P2ry12*^MG-KO^ mice. **h**, Representative images and quantification of P-CEBP/β in Iba1^+^ microglia in the PVN of naive *P2ry12*^fl/fl^ mice and 3-day L-NAME-treated *P2ry12*^fl/fl^ and *P2ry12*^MG-KO^ mice. Arrows indicate the cells containing phosphorylated CEBP/β. Each dot indicates the average of four 160 ×160 μm^2^ FOVs in the PVN of one mouse. **i**, Schematic outline of experiments in (j-l), tamoxifen-treated *Cx3cr1*^CreERT2/+^ mice received bilateral PVN injection of lentivirus which expressed either *Cebpb* shRNA or scramble shRNA under the control of Cre, and 2 wk later, they were subjected to L-NAME challenge. **j**. Representative images and quantification of the ratio of P-C/EBPβ^+^ microglia in the mice treated with lentivirus expressing the indicated shRNA. **k**, PVN microglial density on day 3 post L-NAME challenge. **l**, Systolic arterial pressure of the mice treated with lentivirus expressing the indicated shRNA. n.s., not significant. * P<0.05,** P<0.01; *** P<0.001; **** P<0.0001 by one-way ANOVA in (a, d, h, k) and two-way ANOVA in (f, g, l) with Tukey’s post hoc analysis.

To investigate the causal nexus between ATP-P2Y_12_ signaling and C/EBPβ, we first analyzed the *Cebpb* mRNA expression in BV2 microglial cells *in vitro*. One-hour incubation of BV2 cells with ATP could significantly upregulate the mRNA level of *Cebpb*, and this change could be abolished by co-treatment with PSB0739 (50 μM), a P2Y_12_ inhibitor (Figure 6d). Next, we investigated the correlation between C/EBPβ phosphorylation, an indicator of C/EBPβ activation, and ATP-P2Y_12_ coupling in microglia *in vivo*. To this end, two cohorts of C57BL/6 and *P2ry12*^MG-KO^ mice received ICV injection of either saline or ATP. One hour later, primary microglia were dissociated from those mice for flow cytometry analysis (Figure 6e). Although the microglia derived from both *P2ry12*-intact C57BL/6 mice and *P2ry12*-deprived *P2ry12*^MG-KO^ mice showed an upregulation of C/EBPβ phosphorylation in response to ATP when compared to their genetically matched counterparts from the mice infused with saline, the degree of increase was markedly greater in the *P2ry12*-intact microglia over the *P2ry12*-deprived microglia (10 folds vs. 2.8 folds) (Figure 6f). There was also a significant difference between the ATP-treated groups, as more phosphorylated C/EBPβ (P-C/EBPβ) present in the wild-type microglia over the P2Y_12_-deprived microglia. Next, we exploited the difference in P2Y_12_ expression on microglia derived from tamoxifen-treated *P2ry12*^MG-KO^ mice. we found that ATP ICV infusion resulted in a greater increase of P-C/EBPβ in the remaining P2Y_12_^+^ microglia than the P2Y_12_^−^microglia from the tamoxifen-treated *P2ry12*^MG-KO^ mice (6.7 folds vs. 4.7 folds), accompanied by a remarkable difference in the intensity of P-C/EBPβ signal favoring the former (576,724±27,658 a.u. vs. 46,743.60±17,507 a.u.) (Figure 6g). Since microglia is known to express other purinergic receptors besides P2Y_12_, the mild increase of phosphorylated C/EBPβ by P2Y_12_-null microglia in response to ATP was possibly mediated by other purinergic receptors. *In vivo*, we found that P-C/EBPβ was significantly increased in PVN microglia on day 3 post L-NAME treatment, whereas this rise was suppressed in the *P2ry12*^MG-KO^ mice subjected to the same treatment (Figure 6h). Together, these results strongly indicate that C/EBPβ phosphorylation is mediated, at least partially, by ATP-P2Y_12_ signaling. To further substantiate a role of microglial C/EBPβ in hypertension, we included two cohorts of *Cx3cr1*^CreERT2/+^ mice which received bilateral PVN injection of a lentivirus which expressed a shRNA targeting *Cebpb* or a scramble shRNA under the control of Cre (Figure 6i). The efficiency of this shRNA in knockdown of C/EBPβ expression was verified *in vivo* by immunohistostaining (Figure 6j). After hypertension induction by L-NAME, we monitored the microglia density in the PVN as well as systemic blood pressure. It showed that suppressing C/EBPβ expression in microglia not only restored normal microglia density in the PVN (Figure 6k), but also profoundly dampened blood pressure increases during the hypertensive challenge (Figure 6l). Collectively, these data support that P2Y_12_-C/EBPβ pathway is actively involved in microglial activation relevant to blood pressure increase in experimental hypertension.

## DISCUSSION

Since industrial revolution, prevalence of hypotension due to hypovolemic stress caused by trauma-induced hemorrhage has been dramatically reduced; instead, with prolonged lifespan and excessive intake of salt/calorie, hypertension has been a rampant health issue^76–79^. It has long been noticed that hypertension is associated with neuroinflammation hallmarked by microgliosis in the PVN which mediates an elevated sympathetic outflow, eventually leading to a vicious cycle of neurogenic BP increase. Moreover, microglial activation in hypertension leads to consequential neuronal damages, precipitating the occurrence of neurodegenerative diseases, *e.g.,* Alzheimer’s disease^80, 81^. However, the molecular link between hemodynamic disturbance and microglial activation has long been missing. In this study, by temporospatial analysis of brain microglia in both L-NAME and Ang II hypertensive models, we identified that the microglia in the PVN were an early population, relative to their counterparts in other CNS regions, responding to BP increase. There were marked accumulation of microglia in the PVN with a shift to activation status indicated by both morphological and transcriptomic analysis. Given that hypertension is a hemodynamic disturbance by nature, the blood vessels and the blood cells experience escalated challenges stemming from shear stress after BP elevation, which triggers their release of ATP^57, 82, 83^. Resonating to these observations, we recently revealed that elevated ATP in the plasma is one of the earliest biochemical events occurring under hypertension, which is at least partly responsible for the systemic inflammation associated with hypertension^56^. In alignment, implanted ATP sensor detected more environmental ATP in the PVN after BP increase. We were able to further demonstrate that microglial purinergic P2Y_12_ receptor mediated the effects of excessive extracellular ATP in recruiting microglia to the PVN and the followed sympathetic activation. This conclusion is substantiated by both pharmacological and genetic approaches.

Intriguingly, we found that microgliosis is most prominent in the PVN compared to other nuclei in the early stage of hypertension. Indeed, implanted ATP sensors detected more extracellular ATP in the PVN than in the cortex after a transient pressor stimulation. However, there was no apparent BBB leakage at the early stage of hypertension. Actually, when comparing the trespassing ability of rhodamine B, a fluorescent chemical with molecular size close to ATP, across BBB between multiple brain regions of normotensive naive mice, we observed that the PVN is the most permeable locus in the brain to vasculature-derived small molecules. The mechanism underpinning the specificity of PVN vasculature was unveiled by brain vasculature imaging which showed that the PVN had the densest blood vessel volume with the most complex topology in comparison to brainstem RVLM, motor cortex and MM. This finding resonates with a recent whole-brain blood vessel imaging study which exhibits that the PVN is one of few brain areas possessing the highest density of blood vessels^65^. In addition, PVN vasculature is featured by higher density, more ramification, smaller diameter and less straightness compared to other brain regions (Figure 4d-h). Thus, our data demonstrates that the variation of vasculature structure may predispose some nuclei sensitive, such as the PVN, to hemodynamic disturbance. Given that PVN contains heterogeneous neuronal subpopulations including pre-sympathetic, vasopressin, and cortisol releasing hormone (CRH) neurons^31, 84–86^, it is possible that the topology of PVN vasculature could simultaneously activate PVN pre-sympathetic, AVP and CRH neurons, synergistically regulating body fluid homeostasis for surviving through a stress, *e.g.* hemorrhagic hypotension. Thus, future study is warranted to investigate whether blood-borne bioactive agents could directly affect AVP and CRH release as well.

Microglia have merged as a key player in the regulation of neuronal function throughout the entire lifespan^87–89^. Upon detecting stress-related signals, microglia undergo metabolic and phenotypical changes^90–93^, thereby regulating neuronal behavior by synaptic pruning and circuitry re-wiring during aging and in neurodegenerative diseases^94, 95^. Recent studies unravel that microglia could directly modulate neuronal activity via non-phagocytotic actions^24–27^, implying that microglia have multifaceted impacts on neuronal activity. Our data herein provides a novel insight that microglial phenotype could be altered by hemodynamic stress-elicited ATP release, through which PVN neuronal activity and the downstream sympathetic activity will be secondarily affected. Vasculature-microglia network in the PVN underpins neurogenic hypertension. Given the facts that both hemodynamic disturbance and microglial pro-inflammatory activation are risk factors of neuronal impairment, it is not surprising that neurodegeneration is a complication secondary to hypertension^96–98^.

In mechanism, we demonstrate that microglia activation is at least partially caused by ATP-P2Y_12_ signaling, underscoring a potential therapeutical target for neurogenic hypertension. Indeed, specifically blocking P2Y_12_ receptor by clopidogrel (Plavix) reduces the recurrent heart attack and stroke^99^, both of which has an input from sympathetic outflow. Notably, this antagonist has a molecular weight of 355 Da with hydrophobic property, indicating its potential permeability across the BBB^100^. Thus, it is possible that the therapeutic effects of clopidogrel may be partially achieved by blocking microglial P2Y_12_ signaling. Together, our work warrants a further exploration of the beneficial effects of P2Y_12_ in treating hypertension-associated complications. In conclusion, these results provide novel insights into a microglial role in mediating hemodynamic disturbance of hypertension to influence PVN pre-sympathetic neuronal activity, which is potentially determined by a distinctive characteristic of PVN vasculature.

## ACKNOWLEDGEMENTS

This work was supported by the National Natural Science Foundation of China (31970898, 82170441, 32170894, 82188102); Natural Science Foundation of Zhejiang Province (LZ19C110001, LZ22C080001). MOE Frontier Science Center for Brain Science & Brain-Machine Integration, Zhejiang University. Medical Health Science and Technology Key Project of Zhejiang Provincial Health Commission (WKJ-ZJ-2014); Key Research and Development Project of Zhejiang Provincial Department of Science and Technology (2021C03105). We thank Dr. Long-Jun Wu in Mayo Clinic in the United States generously providing the *P2ry12*^fl/fl^ mice in this study. We acknowledge the technical support by the Core Facility, Zhejiang University School of Medicine. We thank Huang Yingying from the Core Facilities, Zhejiang University School of Medicine for her technical support in primary microglia for bulk RNA-seq.

## AUTHOR CONTRIBUTIONS

B.W. performed the majority of the experiments, analyzed the data and prepared the figures. B.W., Q.S., Q. B., C.L., M.H., N.C., J.H., H.L. and C.L. performed the blood pressure analysis and extracellular ATP detection. G.C., B.W., C.Y. performed the microglial dissociation and RNA-seq data analysis. L.L, performed the LC-MS analysis, and Q. B. and Z.H. performed the electrophysiological recording. X.M. and S.W. provides intellectual input. X.L., X.Z.S. and P.S. designed the experiments. X.Z.S. and P.S. supervised the overall project and wrote the manuscript.

## DECLEARATION OF INTERESTS

The authors declare that no competing financial interests.

## Methods

### Animals

C57BL/6 mice were purchased from Beijing Vital River Laboratory Animal Technology Co., Ltd.. *Cx3cr1*^GFP/GFP^ mice (Stock number: 005582) and *Cx3cr1*^CreERT2/+^ mice (Stock number: 021160) were obtained from Jackson Laboratory (ME, USA). *P2ry12*^fl/fl^ mice were generously provided by Dr. Long-Jun Wu (Mayo Clinic, Rochester, MN, USA). *Cx3cr1*^CreERT2/+^ mice and *P2ry12*^fl/fl^ mice were bred in-house to generate *Cx3cr1*^CreERT2/+^::*P2ry12^f^*^l/fl^ mice. All lines were housed in specific pathogen-free animal facility, and the experimental procedures were approved by Zhejiang University IACUC (ZJU20200007) in accordance with the Guide for Care and Use of the Laboratory Animals.

### Reagents

L-NG-Nitroarginine Methyl Ester (L-NAME, Bachem, China, Catalog #4001301-0025) was fed in the dose of 1.5 mg/ml in drinking water. Angiotensin (Ang) II (GL Biochem, China) was administrated at the dose of 500 ng *per* kg bodyweight (BW) *per* minute via an indwelling osmotic mini-pump (Alzet Model 1002) subcutaneously. Hydralazine (Sigma, USA, Catalog #H1763) was fed in the dose 0.32 mg/ml in drinking water. Apyrase (Sigma, USA, Catalog #A6410) and P2Y_12_ antagonist PSB-0739 (Tocris Bioscience, UK, Catalog #3983) was administered at the dose of 0.1 mg/kg BW/day and 15 μg/day, respectively, through an osmotic mini-pump (Alzet Brain Infusion Kit 3) into the intracerebroventricule (ICV). BrdU (Yeasen, China, Catalog #40204ES60) was injected i.p at the dose 50 mg/kg BW. ATP (MCE, China, Catalog #HY-B2176) was injected intracerebroventricularly at the concentration of 100 mM in a total of 5 μl.

### Surgical procedures

#### Intracranial injection

Animals were anesthetized by inhalation of 1.5∼2% isoflurane mixed with O_2_. The entire procedures were performed on a homeothermic heating pad at 37℃. Mouse head was positioned leveled onto a stereotaxic apparatus (RWD, China, Catalog #68018/68801), followed by drilling a hole at the dimension of 0.5×0.5 mm using a micro-drill (Soeyang, Korea) to gain the injector access into the left cerebral ventricle (A/P: −0.22mm; M/L: 1 mm; D/V:1.6∼1.8 mm).

For parenchymal ATP detection, AAV9-CAG-cATP1.0-WPRE virus (Weizhen Bioscience Inc., China, Catalog #YL006010) was injected to left PVN (A/P: 0.7∼0.9 mm; M/L: 0.2 mm; D/V: 4∼4.2 mm), and left motor cortex (mPFC) (A/P: 1.85 mm; M/L: 0.4 mm; D/V: 1.85 mm) at the titer of 1×10^12^ vg/ml in a total of 150 nl in the aid of nanoliter injector (Nanoject III, Drummond scientific, USA, Catalog #68018).

For *Cebpb* knockdown in microglia, lentivirus rLV-CMV-DIO-shRNA (*Cebpb*)-mEF1a-Pure-WPRE (BrainVTA, China, Catalog #LV-1715) or Lentivirus rLV-CMV-DIO-shRNA (Scramble)-mEF1a-Pure-WPRE (BrainVTA, China, Catalog #LV-1710) was injected to PVN (A/P: 0.7∼0.9 mm; M/L: 0.2 and −-0.2 mm; D/V: 4∼4.2 mm) at the titer of 1×10^9^ TU/ml in a total of 300 nl in the aid of nanoliter injector.

For retrograde labeling RVLM-projecting PVN neurons, mice received ipsilateral RVLM injection (A/P: −6.48∼-7.08 mm; M/L: 0.8∼1.2 mm; D/V: 4.3∼4.7 mm) of retrograde fluorescence microbeads (Lumafluor Inc., Red Retrobeads TM IX; 100∼150 nl).

#### Implantation of optical fiber

To detect extracellular ATP signal in the PVN and cortex, two optical fibers were implanted using the following coordinates: PVN: A/P: −0.63 mm, M/L: 0.2 mm, D/V: 4 mm; and mPFC: A/P: 1.85 mm; M/L: 0.4 mm; D/V: 1.85 mm. Fibers were stabilized on the surface of the skull by dental cement.

### Vascular permeability and tracing assessments

For vascular permeability analysis, mice were anesthetized by inhalation of 1.5∼2% isoflurane. Rhodamine B (354 μg/g BW; 479Da MW, Ex/Em: 553/627, Aladdin, China, Catalog #A104960), 10 kDa FITC-dextran (15 μg/ g BW; Ex/Em: 494/520 nm, Sigma, USA, Catalog #FD10S) or 70 kDa FITC-dextran (100 μg/g BW; Ex/Em: 494/520 nm, Sigma, USA, Catalog #FD70S) were i.v. injected. Fifteen minutes after Rhodamine B infusion or 4 hr after FITC-dextran infusion, transcardiac perfusion was performed. Brains were then dissected in the aid of brain matrix (RWD, China, Catalog #68707), followed by homogenization in 5 μl/mg tissue PBS+0.1% Triton X-100 (PBST). After centrifuging at 12,000 g for 20 min at 4℃, the supernatant was collected for fluorescence intensity analysis by a microplate reader (Agilent, BioTek Synergy H1).

For tracing analysis, animals were deeply anesthetized with isoflurane. The 150 kDa FITC-dextran (Sigma, USA, Catalog #46946) was first dissolved in 0.9% saline, and was i.v. administrated at the dose of 100 mg/kg BW. Five minutes post injection, mice were euthanatized and the brains were isolated and fixed in 4% PFA overnight at 4℃. After that, the samples were immersed in 30% sucrose for 72 hr before being sectioned into 100 μm coronary slices for vasculature reconstruction, or into 30 μm slices for immunohistostaining.

### Fiber photometry *in vivo* recording of ATP signaling

To monitor the real-time ATP sparkling in the targeted nuclei, a multi-channel fiber photometry system (Inper, China, Catalog #SPP0103) was used. Two weeks post optical fiber implantation, recording system was connected to the fibers. ATP signals in the tissues were recorded in the wave length range of 500∼540 nm under the excitation wave length of 450∼490 nm. A reference channel (excitation wave length of 390∼430 nm, emission wave length of 480∼560 nm) was used to normalize movement-induced background noise. On recording, 30 min-baseline signals were first collected at a frequency of 30 Hz; an i.p. injection of Ang II (7.2 mg/kg BW) was followed; afterwards, the signals in another 30-min duration were recorded. The florescent signals simultaneously recorded from the PVN and mPFC in the same animal were analyzed using the formula (F_x_-F_Baseline_)/F_Baseline_ × 100%.

### Hypothalamic brain slice preparation for electrophysiological recording

Mice were anesthetized by isoflurane and decapitated. Brains were cut into 300 mm coronal slices containing the hypothalamic PVN using a vibratome (Leica VT1200S) in ice-cold cutting solution saturated with 95% O2 and 5% CO2, containing (in mM): 200 sucrose; 2.5 KCl; 1.25 NaH2PO4; 25 NaHCO3; 0.5 CaCl2; 7 MgCl2; 25 D-glucose; 0.6 sodium pyruvate; 1.3 ascorbic acid. Sliced sections were then incubated in solution containing (in mM): 110 sucrose; 2.5 KCl; 1.25 NaH2PO4; 25 NaHCO3; 0.5 CaCl2; 7 MgCl2; 25 D-glucose; 0.6 sodium pyruvate, 60 NaCl; 1.3 ascorbic acid at 32℃ for 20 min before recording.

### Electrophysiological recordings

Slices were transferred into a recording chamber perfused with artificial cerebrospinal fluid (ACSF) containing (in mM): 15 sucrose; 2.5 KCl; 1.25 NaH2PO4; 25 NaHCO3, 2 CaCl2; 1.2 MgCl2; 25 D-glucose; 0.6 sodium pyruvate; 125 NaCl; 1.3 ascorbic acid, and bubbled with 95% O2-5% CO2. Fluorescent labeled (retrograde microbeads injected to RVLM) neurons in the PVN were identified under a fluorescence microscope (Olympus BX51) for whole-cell patch-clamp recordings. Microelectrodes (6–8 MU) were pulled from borosilicate glass and filled with the intracellular solution (pH 7.4, 295 mOsm). Whole-cell patch clamp recordings were performed using an Axopatch 700B amplifier (Axon Instruments) filtered at 2 kHz. The membrane resistance, capacitance and spontaneous firing were recorded under current-clamp mode without current injection. To test the PVN neuronal excitability and spike-frequency adaptation, 300-ms depolarizing currents pulses with different intensities were injected to the clamped cells and the evoked action potential was recorded.

### Systemic arterial pressure recording

#### Telemetry recording

Systolic arterial pressure was recorded via the indwelling blood pressure telemetry transmitter (DSI, USA, HD-X11); data were collected, stored and analyzed using Dataquest A.R.T. 4.0 software (Data Science International) at 60 seconds per 5 minutes for at least 1hour between 10am-4pm.

#### Tailcuff recording

Systolic arterial pressure was determined in conscious mice using a computerized non-invasive tail cuff system (Visitech systems, Apex, BP-2000 series II) as previously reported^24, 47, 101^. Each mouse was trained for 5 consecutive days prior to data acquisition. Systolic arterial pressure was determined by averaging 25 measurements. The tracing waves were manually reviewed to verify proper blood pressure determination.

#### Acute systemic arterial pressure recording

Acute systemic arterial pressure was determined in the mice anesthetized by inhalation of 1.5% isoflurane mixed with O_2_. The right common carotid artery was cannulated for arterial blood pressure recording. Data were sampled via a pressure transducer (AD Instruments; Catalog #MLT0699), and the signals were digitized and analyzed at rate of 1,000 Hz using PowerLab acquisition system (AD Instruments v8.0).

### Liquid chromatographic mass spectrometry (LC-MS)

Plasma NE and ATP were quantitatively analyzed by LC-MS (Agilent 1260; Agilent Technologies, USA) coupled with API4000 mass spectrometer (AB Sciex, USA) in positive electrospray ionization (ESI) through multiple reaction monitoring (MRM) mode. Data were acquired by Analyst software Hotfixes (AB Sciex, Ontario, Canada).

### Immunohistochemistry

After transcardiac perfusion, brains were collected and fixed in 4% PFA at 4℃ overnight, followed by being immersed in 30% sucrose for 72 hr. Samples were sectioned by a cryostat at −20°C (Leica, Germany, SM2010 R), and 30 μm coronal slices were made. Floating brain slices were blocked with 5% donkey serum in 0.2% Triton X-100 (Sigma, USA, X100) PBS for 3 hr at room temperature, followed by incubation with primary antibodies at 4℃ overnight, and then with corresponding secondary antibodies for 2 hr at room temperature. Brain sections were mounted onto glass slides covered by DAPI mounting medium (Southern Biotech, USA, 0100-20).

The following primary antibodies against murine antigens were used: rabbit (1:1000. Wako, Japan, Catalog #019-19741) or goat (1:1000. Wako, Japan, Catalog #011-27991) anti-Iba-1, rabbit anti-phospho-CEBP/β (1:500, Invitrogen, USA, Catalog #PA5-104823), rabbit anti-α-SMA (1:1000, Abcam, UK, Catalog #ab5694), rat anti-CD13 (1:1000, Abcam, UK, Catalog #ab33489), rabbit anti-Ki67 (1:500. Servicebio, China, Catalog #GB111499). In addition, an Alexa488-conjugated anti-BrdU antibody (1:500, Biolegend, USA, Catalog #364105) was used.

The secondary antibodies were all purchased from Abcam UK and applied with 1:500 dilution. Donkey anti-rabbit (Catalog #ab150075, #ab175470), donkey anti-goat (Catalog #ab150131) and donkey anti-rat (Catalog #ab150155) antibodies were selectively used.

### Morphological analysis

All images were captured by confocal microscope (ZEISS LSM 900) and analyzed by Imaris software (v9.5). We used the following coordinates for microglia and vasculature analysis: PVN as A/P: −0.6∼-1.2 mm; M/L: 0.1∼0.75 mm; D/V: 4.25∼4.75 mm; RVLM as A/P: −6.48∼-7.08 mm; M/L: 0.8∼1.2 mm; D/V: 4.3∼4.7 mm; cortex A/P: −0.6∼-1.2 mm; M/L: 2.5∼3.5 mm; D/V: 0.2∼1.5 mm.

#### Microglial morphological analysis

Iba1^+^ microglial processes were traced and analyzed using the “Filament Component” module of Imaris in z-stacks of 3D reconstructed images. Microglial soma was determined manually by a spherical tracing. All the processes stemmed from the center of the sphere were analyzed manually. Sholl analysis was performed by computing interaction number of the traced processes in concentric circles created from the center of the sphere which indicate the microglial soma. The interval between adjacent concentric circle is 1 μm. The counts of the interaction indicate the complexity of microglial processes. Soma volume of individual microglia was computed based on the diameter of the traced sphere.

#### Microglia density analysis

Microglial (soma diameter ≥8 μm) density was analyzed by “Spots Component” module of Imaris.

#### Cerebral vasculature analysis

Vasculatures in the PVN, RVLM, S1BF and MM were traced with “Filament component” module of Imaris to indicate the levels of straightness, length and branch points. To further unravel straightness of the vasculature, a rainbow colors were coded according to the straightness analysis. Volumes of vessels in above regions were quantified according to the reconstruction using the “Surface Component” module of Imaris.

### Microglia dissociation

Dissected brain tissues from entire cortex, hypothalamus and brainstem (2×2×2 mm^3^ dimension for brain regions including the latter two nuclei) were minced with blade on ice individually, followed by enzymatic digestion with 0.6 mg/ml collagenase IV (Worthington, USA, Catalog #LS004189) and 100unit DNase I (Sigma, USA, Catalog #DN25) in 6 ml DMEM for each brain at 37°for 40 min. The dissociated tissues were resuspended in cold PBS after centrifugation (300 g for 5 minutes at 4℃) and filtered (70 μm cell strainer, biologix, USA, Catalog #15-1070). Microglia from the filtrate were enriched by centrifugation in discontinuous 37% and 70% Percoll (GE, USA, Catalog #17-0891-09).

For the sample for microglia sorting, brain tissues from *Cx3cr1*^GFP/GFP^ mouse were added as carrier in the samples to protect microglia loss during the entire dissociation procedure^102^. In the flow cytometry analysis, the carrier microglia could be recognized and excluded by their GFP expression.

### Flow cytometry analysis

Dissociated single cells were resuspended and washed twice with 200 μl FACS buffer (Ca^2+^ and Mg^2+^ free PBS with 2.5mM EDTA and 0.1-1% BSA, 0.1% NaN_3_). After blocking Fc receptors (anti-CD16/32 antibody, Biolegend, USA, Catalog #101320) at 4℃ for 10 minutes, cells were subject to surface staining by being incubated with a cocktail of anti-CD11b (Biolegend, USA, Catalog #101225), anti-CD45 (Biolegend, USA, Catalog #103126) and anti-P2Y_12_ (Biolegend, USA, Catalog #848003) antibodies at 4℃ for 20 minutes.

To stain intracellular phosphorylated C/EBPβ, cells were first incubated with a rabbit anti-mouse C/EBP β primary antibody (Invitrogen, Catalog #PA5-104823) which specifically targets on the phosphorylated Thr188 of murine C/EBPβ (a conservative locus to Thr235 of human C/EBPβ) in eBioscience™ Transcription Factor Staining Buffer Set (Invitrogen, Catalog #00-5523-00). For background signal staining, a rabbit IgG isotype antibody (Invitrogen, Catalog #02-6102) was used as the primary antibody. Cells were then incubated with a fluorophore-conjugated donkey anti-rabbit IgG secondary antibody, followed by flow cytometry (Agilent Novocyte) analysis. Data were processed with FlowJo (v10).

### Fluorescence-activated flow cytometry sorting

Dissociated single cells were resuspended and washed twice with 200 μl FACS buffer (Ca^2+^ and Mg^2+^ free PBS with 2.5 mM EDTA and 0.1-1% BSA, 0.1% NaN_3_). After blocking Fc receptor by CD16/32 antibody (Biolegend, USA, Catalog #101320) at 4℃ for 10 minutes, cells were incubated with a cocktail of CD11b-APC-Cy7 (Biolegend, USA, Catalog #101225) and CD45-pacific blue (Biolegend, USA, Catalog #103126) antibodies at 4℃ for 20 minutes. Cells were also incubated with 7-AAD (Biolegend, USA, Catalog #420404) prior to sorting. Microglia were sorted based on their CD11b^+^/CD45^low^/7-AAD^-^/GFP^-^ profile. In general, we obtained 2,169±854.945 microglia cells from hypothalamus, 2,125.33±852.34 from brain stem, and 10,011.67±11.67 from cortex.

### Bulk RNA-seq library construction

We utilized a recently developed method Holo-Seq, which is applicable to low cell number samples^24, 103^, to carry out the RNA-seq library construction with limited number of microglia cells or macrophages after FACS. Holo-Seq introduces artificial carrier RNA to the library construction procedures, which protects trace amount of sample RNA from loss during library construction procedures, and adapts the standard bulk RNA-seq method to our low-cell-number samples, achieving high quantification accuracy and detection sensitivity of the whole transcriptome profiling. In brief, samples obtained by FACS together with 100 ng carrier RNA (PolyA- and PolyA+ mixed in 98:2 ratio) were lysed by Buffer RLT Plus (Qiagen, USA). Then we used PolyA mRNA Magnetic Isolation Module (NEBNext, USA, Catalog #E7490) to obtain PolyA-tailing RNAs, which were then presented to the common bulk RNA-seq library construction procedures, following the protocol of Ultra RNA Library Prep Kit for Illumina (NEBNext, USA, Catalog #E7530). The carrier RNAs were around 200 nt and transcribed in vitro, DNA templates of which have Not I restriction enzyme cutting sites every 20-30 bp. cDNAs reversely transcribed from carrier RNAs were removed from the RNA-seq library by Not I (NEB, USA, Catalog #R0189L) digestion after the adaptor ligation step, so that the final sequencing result could not be affected by the carrier RNAs. Libraries passed the Agilent 2200 Bioanalyzer (Agilent) quality check were sent for sequencing on the Illumina HiSeq X Ten platform (150 bp, paired-end mode).

DNA template for carrier RNA in vitro transcription (PolyA-carrier RNA, 233 bp) TAATACGACTCACTATAGGGCGGCCGCATATTAACGCTTACAATTTAGCGGCCGCGTGGCACTTTTCGGGGAAATGCGGCCGCTGCGCGGAACCCCTATTTGTTGCGGCCGCTATTTTTCTAAATACATTCAAGCGGCCGCATATGTATCCGCTCATGAGACGCGGCCGCAATAACCCTGATAAATGCTTCGCGGCCGCTAATATTGAAAAAGGAAGAGCGGCCGCTATGA

DNA template for carrier RNA in vitro transcription (PolyA+ carrier RNA, 229 bp) TAATACGACTCACTATAGGGCGGCCGCATATTAACGCTTACAATTTAGCGGCCGCGTGGCACTTTTCGGGGAAATGCGGCCGCTGCGCGGAACCCCTATTTGTTGCGGCCGCTATTTTTCTAAATACATTCAAGCGGCCGCATATGTATCCGCTCATGAGACGCGGCCGCAATAACCCTGATAAATGCTTCGCGGCCGCAAAAAAAAAAAAAAAAAAAA AAAAAAAAAAAA

After sequencing and alignment, we obtained 16.9±3.0 M transcriptome mapped reads for microglia derived from the hypothalamus (n=9), 16.1 M ± 3.3 M transcriptome mapped reads for the brain stem (n=9) and 14.1±2.1 M transcriptome mapped reads for the cortex (n=9). After analysis, there were 14573±800 genes for microglia derived from the hypothalamus (n=9), 15136±911 genes for the brain stem (n=9) and 15758 ±573 genes for the cortex (n=9).

### RNA-seq data quality control and quantification

We used BBDuk (BBMap v 38.86) to trim adaptor and low-quality bases off the raw sequencing reads, during which process Illumina adaptors and sequence regions with average quality score bellow 15 were removed. Reads shorter than 36 bp were discarded after trimming. Then we aligned the left reads to the mouse genome (GRCm38, Genecode) by STAR (v2.7.5a_2020-6-19) with default parameters in paired-end mode, and quantified the transcript abundance with the ‘quant’ step of Salmon (v1.2.1).

### Bulk RNA-seq analysis

With the gene mapped read count data obtained after quantification, we performed differential gene expression analysis in microglia from naive mice or Ang II or L-NAME treated mice, respectively in 3 different brain sections. Firstly, we filtered out genes those weren’t expressed (weren’t expressed in at least two out of three samples) in neither naive nor hypertension-induction group, then we did gene differential expression analysis via R package edgeR (v3.28.1) with function exactTest. We defined differential expressed genes (log2FC>1, or log2FC<-2 with P_value_ < 0.05) and found the common DEGs after Ang II and L-NAME treatments. Then we used the common DEGs to do the gene ontology (GO) analysis with the enrichGO function of package clusterProfiler (v4.0.5). Besides, with the common DEGs (overlapping DEGs in Ang II and L-NAME treatment) of three different brain regions (hypothalamus, brain stem and cortex), we obtained region-specific DEGs of each region by excluded DEGs appearing in other two regions for further observations.

For hierarchical clustering analysis of samples, we firstly obtained their Euclidean distance matrix by R function dist with the gene TPM values of each sample, then we used the agglomeration method of ‘complete’ to do the hierarchical clustering by R function hclust.

KEGG pathway KGML files and pathway related information was downloaded with KEGGREST^104^, then parsed to data.frame. *P2ry12* downstream signal analysis was conducted with igraph^105^, and visualized with ggraph^106^ and tidygraph^107^

### Cell culture

Murine microglial cell line BV2 was cultured in high glucose DMEM medium (Corning, USA, Catalog #10-013-CV) with 10% FBS (Bovogen, Australia, Catalog #SFBS-AU), 10 mM HEPES (Gibco, USA, Catalog #15630080), 100 U/ml penicillin and 100 μg/ml Streptomycin (BBI, China, Catalog #E607018). BV2 cells were cultured in the presence of 5 mM ATP (MCE, China, Catalog #HY-B2176) alone, or together with 50 μM PSB-0739 in the DMEM medium supplemented with 1% FBS for 1 hr.

### Real-time RT-PCR

Total RNA extraction was performed by using RNA purification kits (ES science, China, Catalog #RN001 for BV2 cells, and Catalog #RN002plus for tissue). Extracted RNA was reversed to cDNA with cDNA Synthesis Kit (Yeasen, China, Catalog #1141ES10/60). Target gene expression was quantified by SYBR Green (Yeasen, China, Catalog #11201ES03/08/60) using corresponding primer mixers as the following:

#### *Gapdh* primers

Forward: AGGTCGGTGTGAACGGATTTG; Reverse: TGTAGACCATGTAGTTGAGGTCA

#### *Pry12* primers

Forward: ATGGATATGCCTGGTGTCAACA; Reverse: AGCAATGGGAAGAGAACCTGG

#### *Cebpb* primers

Forward: CAACCTGGAGACGCAGCACAAG; Reverse: GCTTGAACAAGTTCCGCAGGGT

Acquired data was normalized to the data of housekeeping gene *Gapdh,* and analyzed to get ΔΔCt. All procedures were performed according to the manufacturer’s instructions.

### Statistics

Data were analyzed by paired-or unpaired t-tests; One-or two-way ANOVA with post hoc tests using Graphpad Prism (v6.0). Statistical significance was considered when P<0.05. Data were shown as mean ± s.e.m.. In the GO term analysis in Figures 2C, R package clusterProfiler (v4.2.2) was used, and the enrichment p-value was calculated with hypergeometric test.

## Data availability

The raw RNA-seq data generated and analyzed in this paper have been deposited in the Genome Sequence Archive^108^ in National Genomics Data Center^109^, China National Center for Bioinformation/Beijing Institute of Genomics, Chinese Academy of Sciences (GSA: CRA008127) that are publicly accessible at https://ngdc.cncb.ac.cn/gsa. Source data are provided with this paper.

## Code availability

This manuscript does not report original code.

---

**Extended Data Figure 1.**
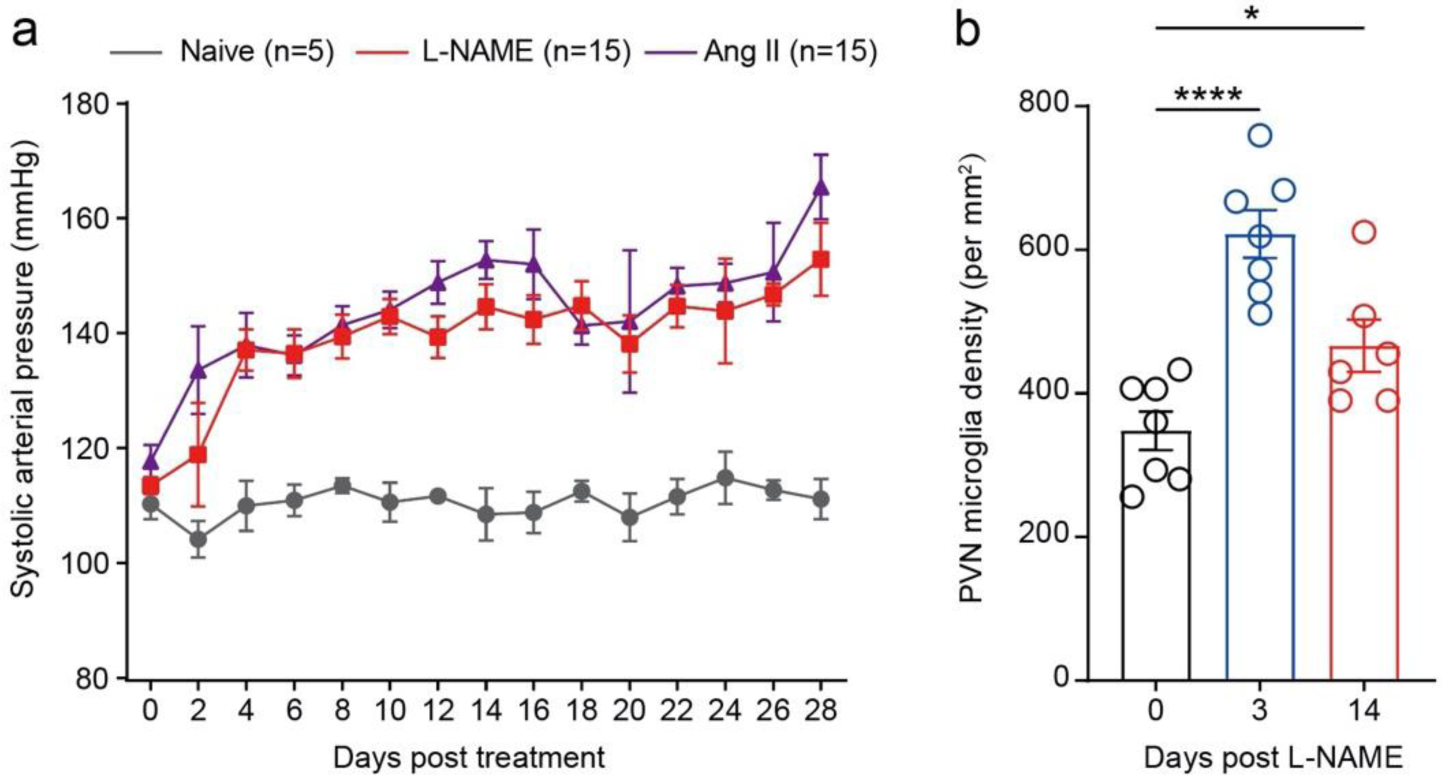
The dynamics of BP increase and microglial density in hypertension models. **a**, Systolic blood pressure of naïve mice or mice treated with Ang II (500 ng/kg/min by subcutaneous minipump) or L-NAME (1.5 mg/ml in drinking water). **b**, Microglia density were analyzed in the PVN in the indicated time points after L-NAME treatment. * P<0.05, **** P<0.0001 by one-way ANOVA with Tukey’s post hoc test.

**Extended Data Figure 2.**
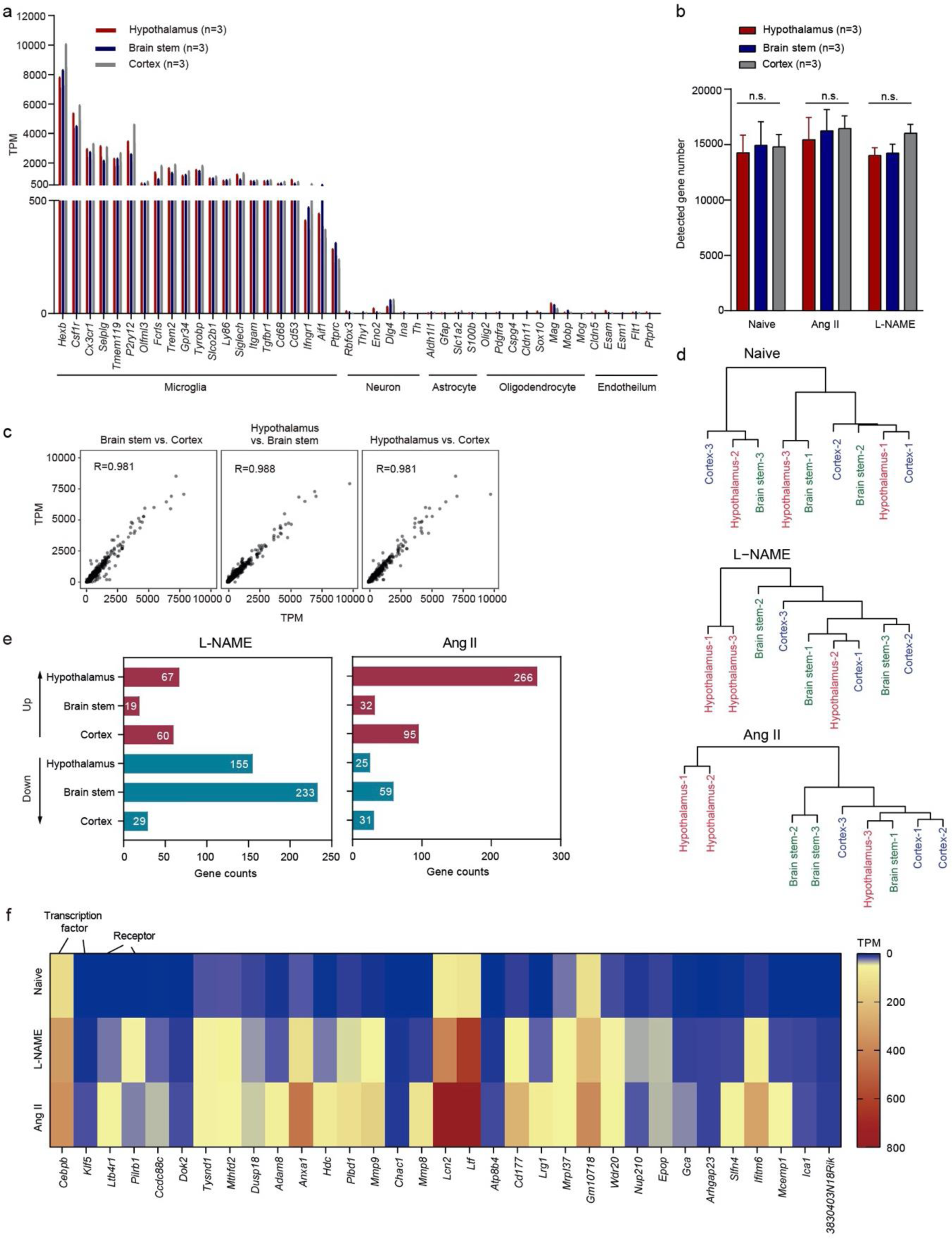
Transcriptomic analysis of microglia derived from normotensive and hypertensive animals. **a**, Normalized expression (TPM) of marker genes for microglia, neuron, astrocyte, oligodendrocyte and endothelial cell in the enriched microglial cells dissociated from hypothalamus, brain stem and cortex of normotensive C57BL/6 mice. **b**, Numbers of detected genes in microglia dissociated from the indicated brain regions derived from the mice with or without 3-day Ang II or L-NAME treatment. **c**, Pearson correlation between bulk RNA-seq data of microglia dissociated from different brain regions of normotensive mice. **d**, Dendrogram indicating hierarchical clustering of bulk RNA-seq samples from the 3 indicated brain regions of normotensive mice and the mice after 3-day treatment of L-NAME or Ang II. **e**, Numbers of differentially expressed genes (DEGs) of microglia enriched from the indicated regions of L-NAME-or Ang II-treated mice compared to the normotensive control mice. **f**, The 33 common upregulated DEGs in hypothalamic microglia dissociated from Ang II and L-NAME hypertensive models.

**Extended Data Figure 3.**
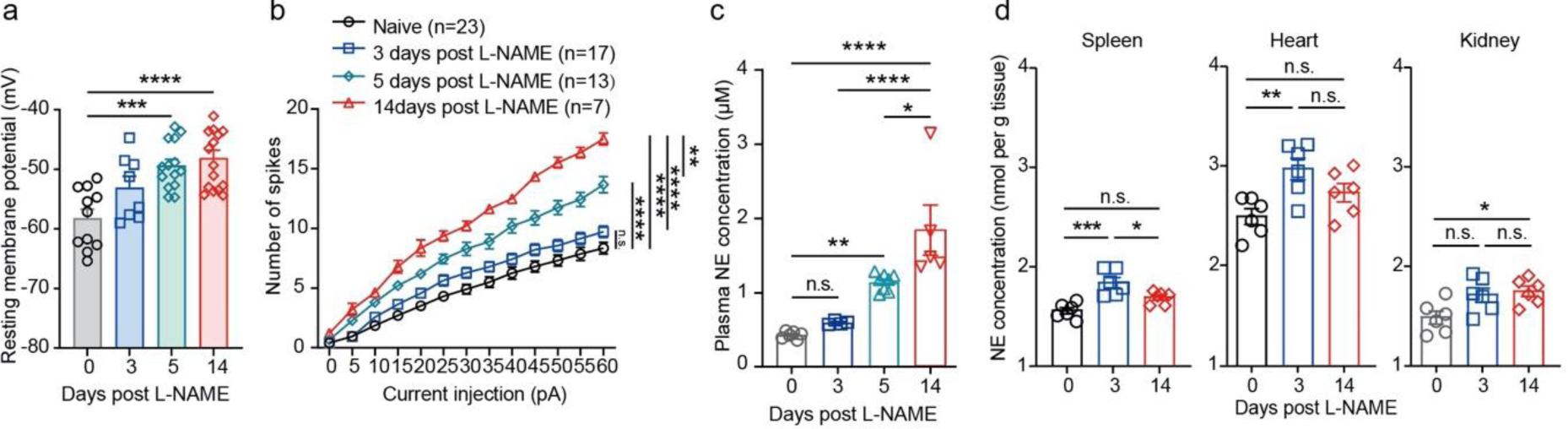
PVN pre-sympathetic neuronal activity and peripheral NE levels during hypertension. **a**-**b**, Resting membrane potential (a) and evoked action potential (b) of RVLM-projecting PVN neurons from C57BL/6 mice post L-NAME treatment. **c-d**, Norepinephrine (NE) levels in the blood plasma (c) and the indicated organs (d) were measured at different time points post L-NAME treatment. * P<0.05, ** P<0.01, *** P<0.001, **** P<0.001 by one-way ANOVA in (a, c, d) and two-way ANOVA in (b) with Tukey’s post hoc analysis.

**Extended Data Figure 4.**
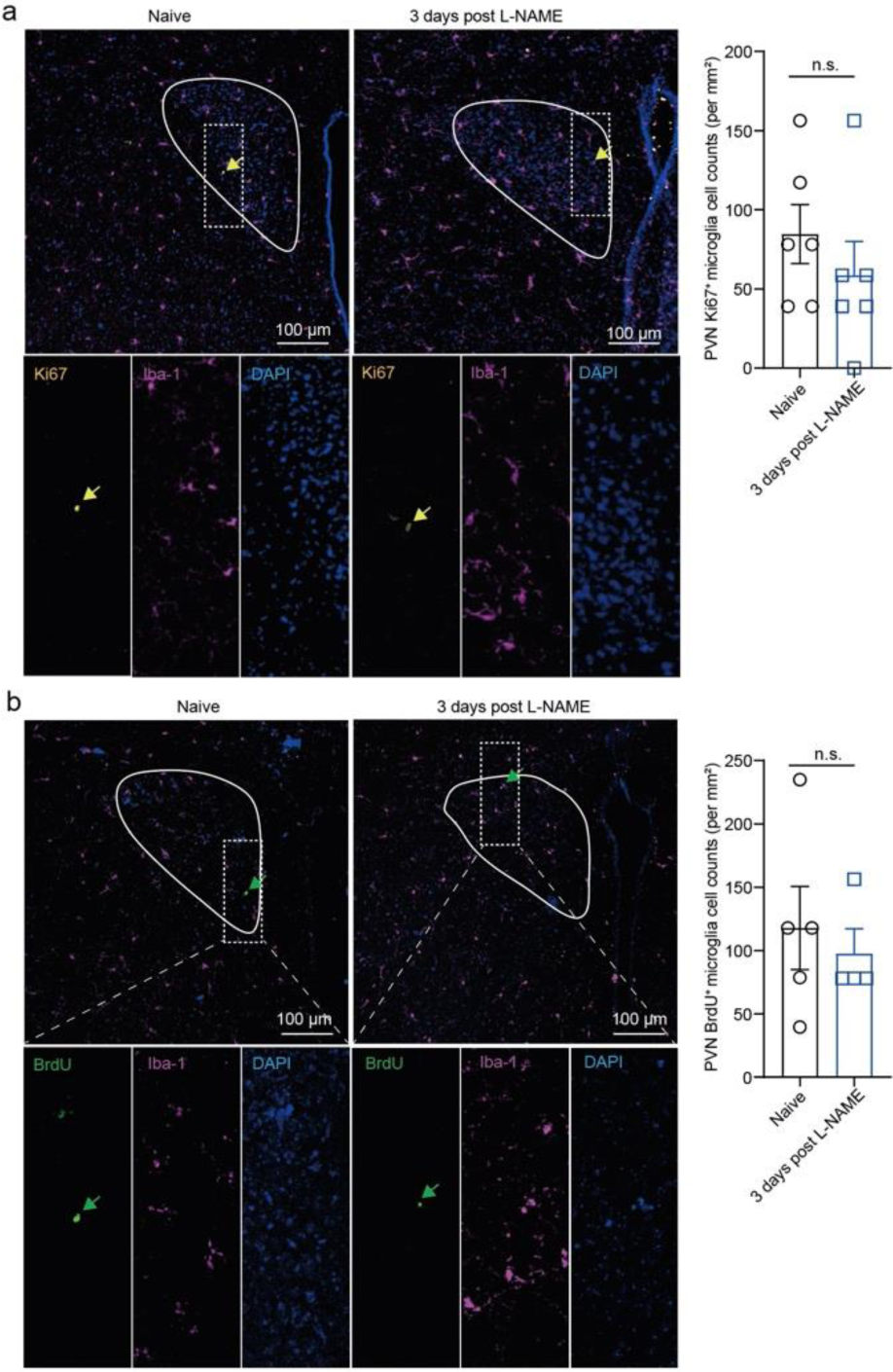
Proliferation analysis of microglia in the early stage of hypertension. Representative images and quantification of co-staining of Ki67 (**a**) or BrdU (**b**) with Iba1in the PVN from the naïve mice and the mice 3 days post L-NAME treatment. For b, mice were i.v. injected with BrdU at 2 hr before brain harvesting. Arrow, Ki67^+^ or BrdU^+^ microglia. The curved lines delineate the contour of PVN. Each dot indicates the average of four 160 ×160 μm^2^ FOVs in the PVN of one mouse. n.s. not significant by unpaired two-tailed t-test.

**Extended Data Figure 5.**
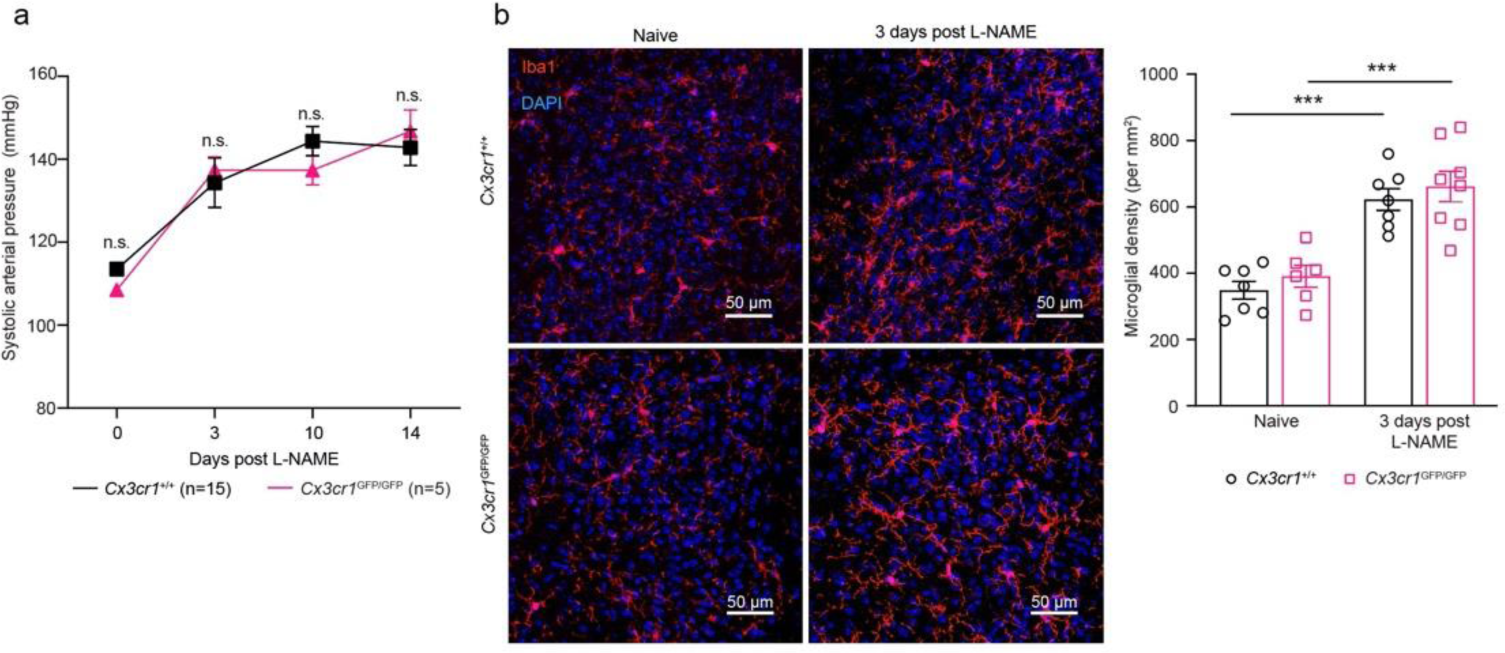
CX3CR1 does not mediate the microglial accumulation in the PVN in the early stage of hypertension and has no effect on the L-NAME-elicited pressor response. **a**, Systolic arterial pressure during L-NAME treatment in *Cx3cr1*^+/+^ and *Cx3cr1*^GFP/GFP^ mice. **b**, Representative images and analysis of microglial density in the PVN of the indicated mice 3 days post L-NAME treated mice. Each dot indicates the average of four 160 ×160 μm^2^ FOVs in the PVN of one mouse. n.s., not significant. *** P<0.001 by two-way ANOVA with Tukey’s post-hoc test.

**Extended Data Figure 6.**
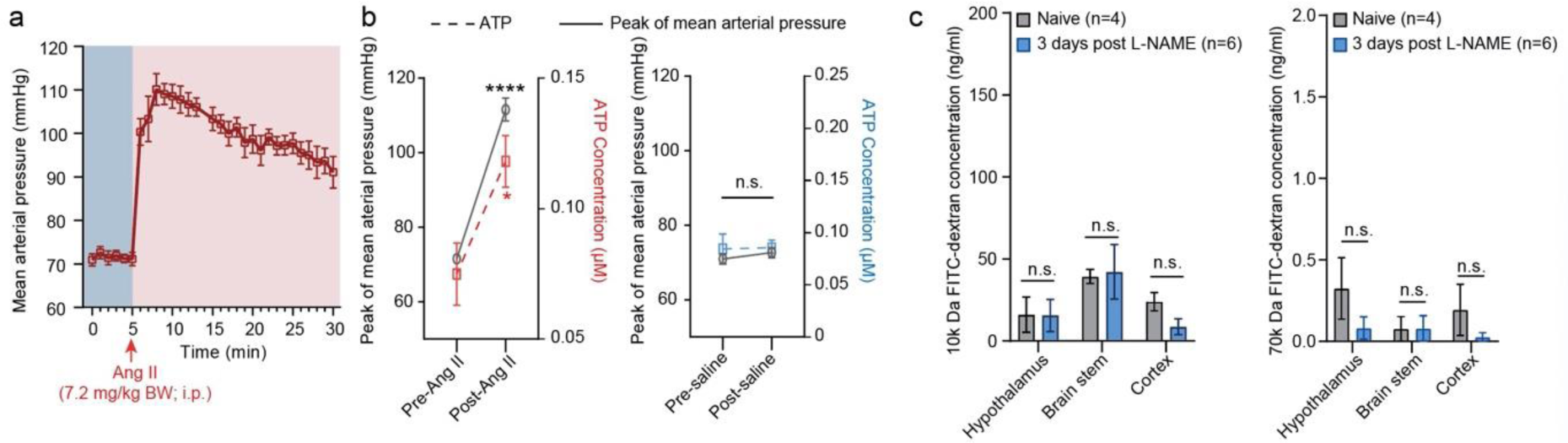
Pressor effects on plasma ATP and BBB permeability. **a**, Under anesthesia, mice received an i.p. injection of Ang II (7.2 mg/kg body weight). The mean arterial BP was measured by an indwelling catheter in the common carotid artery (n=6). **b**, Effects of a bolus i.p. injection of either Ang II (left) or saline (right) on systolic arterial pressure and plasma ATP in C57BL/6 mice 2 min post either Ang II or saline treatment. n = 6 for BP recording, n = 5 for plasma ATP analysis. **c**, Fifteen minutes after i.v. injection of 10 kDa or 70 kDa FITC-dextran, mice were transcardiacally perfused with PBS, and the FITC-dextran in the parenchyma of the indicated brain regions were measured. n.s. not significant, **** P<0.0001 vs. baseline by paired t-test in (b).

**Extended Data Figure 7.**
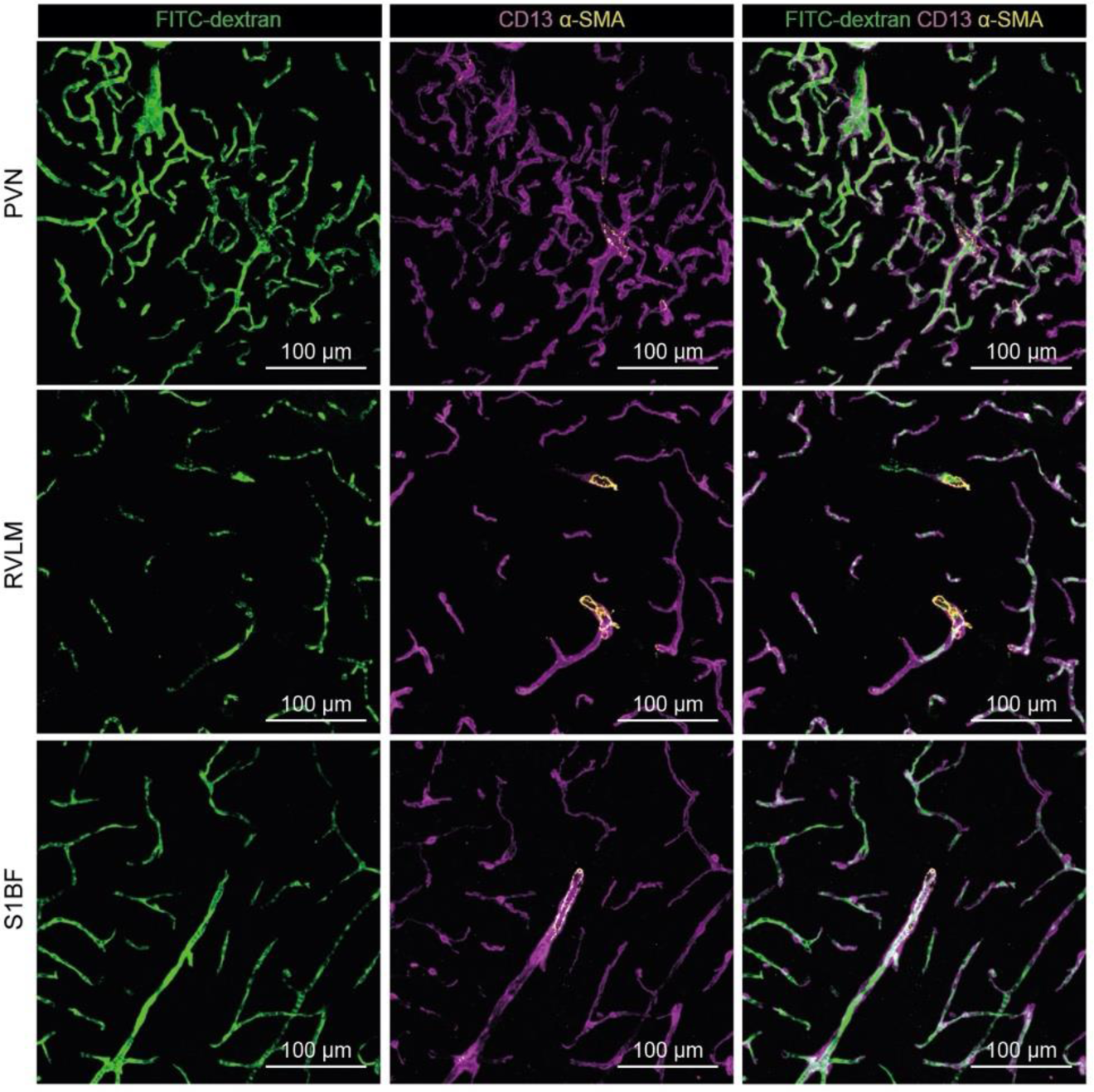
Illustrating the cerebral vasculature. Representative images of FITC-dextran-labeled vessels, CD13^+^ pericytes and α-SMA^+^ vascular smooth muscle cells in the indicated nuclei of naïve mice.

**Extended Data Figure 8.**
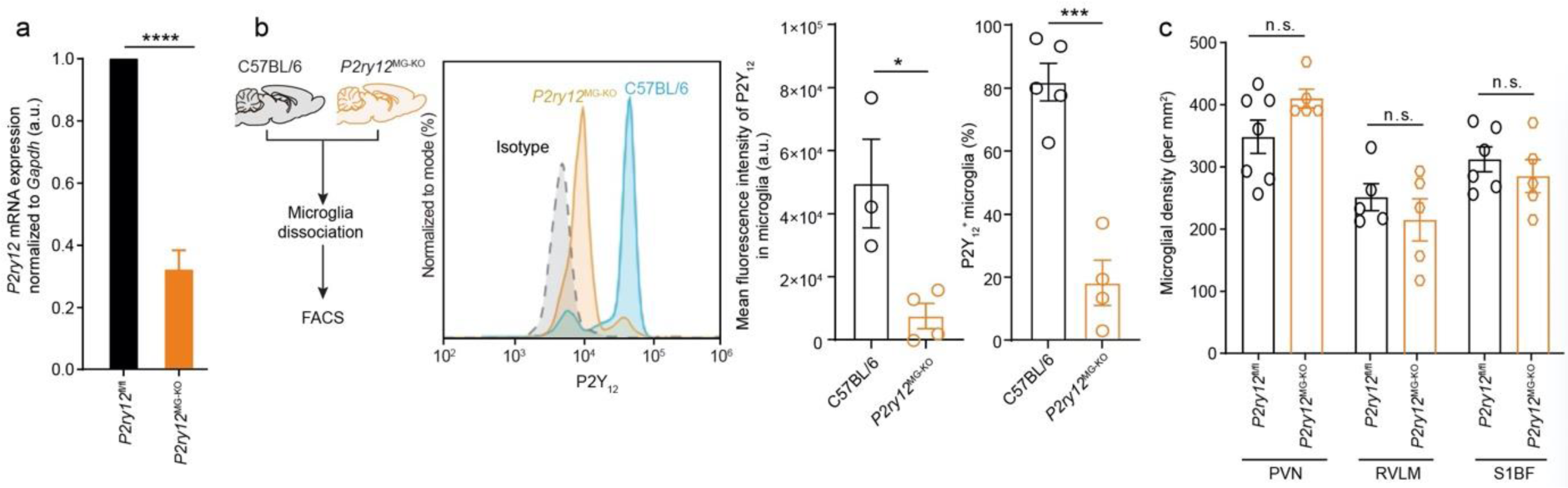
Efficacy of *P2ry12* ablation and effects on microglial density. **a**, Real-time RT-PCR analysis of *P2ry12* mRNA expression in the brain tissues dissociated from *P2ry12*^fl/fl^ mice or *P2ry12*^MG-KO^ mice 3 weeks post tamoxifen treatment. n = 4. **b**, Experimental protocol for P2Y_12_ expression analysis in C57BL/6 and *P2ry12*^MG-KO^ mice (left panel). CD11b^+^CD45^low^ microglia were pre-gated and the representative histogram showing their P2Y_12_ expression in above two cohorts of mice (middle panel). Quantification of microglial P2Y_12_ expression in mean fluorescence intensity and in percentage of P2Y_12_ ^+^ cells was also shown (right panel). **c**, Microglial density analysis in the PVN, RVLM and S1BF of *P2ry12*^fl/fl^ or *P2ry12*^MG-KO^ mice 3 weeks post tamoxifen treatment. n.s., not significant. * P<0.05, *** P<0.001, ****P<0.0001 by unpaired t-test.

